# Distinct representations of innate and learned threats within the thalamic-amygdala pathway

**DOI:** 10.1101/2023.01.10.523445

**Authors:** Valentina Khalil, Islam Faress, Noëmie Mermet-Joret, Peter Kerwin, Keisuke Yonehara, Sadegh Nabavi

## Abstract

Behavioral flexibility and timely reactions to salient stimuli are essential for survival. The subcortical thalamic-basolateral amygdala (BLA) pathway serves as a shortcut for salient stimuli ensuring rapid processing. Here, we show that BLA neuronal and thalamic axonal activity mirror the defensive behavior evoked by an innate visual threat as well as an auditory learned threat. Importantly, perturbing this pathway compromises defensive responses to both forms of threats, in that animals fail to switch from exploratory to defensive behavior. Despite the shared pathway between the two forms of threat processing, we observed noticeable differences. Blocking beta-adrenergic receptors impair the defensive response to the innate but not the learned threats. This reduced defensive response, surprisingly, is reflected in the suppression of the activity exclusively in the BLA, as the thalamic input response remains intact. Our side-by-side examination highlights the similarities and differences between innate and learned threat-processing, thus providing new fundamental insights.

---

Survival is the direct product of maximizing gains and avoiding harms. This is achieved by integrating brain circuitry that ensures survival through environment exploration while maintaining threat detection and avoidance (Blanchard et al., 2001; Evans et al., 2019; Gross & Canteras, 2012; Headley et al., 2019; Orsini & Maren, 2012; Silva et al., 2016). Such capacity can be acquired through learning, a mechanism widely believed to rely on synaptic plasticity (Mongeau et al., 2003; Johansen et al., 2011; Quirk et al., 1995; Rogan et al., 1997; Tierney, 1986). However, specific sensory stimuli could be innately appetitive or aversive and evoke approach or defensive behaviors, respectively (Pereira & Moita, 2016).

Rapid and continuous integration of innate and learned mechanisms promotes survival. This integration is ideally carried out as a timely and appropriate reaction to sensory stimuli. While cortical processing is necessary for cognitively demanding tasks, it is not well suited for rapid threat detection and response. However, subcortical processing serves as a neural shortcut that can crudely and rapidly elicit defensive behaviors, bypassing the more deliberate and intricate cortical processing. This efficient processing is partially explained by the fact that subcortical areas are among the earliest brain areas that receive sensory information (Carr, 2015; McFadyen et al., 2020; Pessoa, 2008; Pessoa & Adolphs, 2010).

It is well established that the subcortical circuit from the multisensory thalamus, medial geniculate nucleus (MGN) to the basolateral amygdala (BLA) is necessary in the acquisition and processing of auditory learned threat conditioning in rodents (Barsy et al., 2020; Edeline & Weinberger, 1992; Iwata et al., 1986; Lee et al., 2021; LeDoux et al., 1984; Romanski & LeDoux, 1992a; Romanski & LeDoux, 1992b; Taylor et al., 2021). However, the role of the subcortical MGN-BLA pathway is underemphasized and understudied in processing innate threats (Kang et al., 2022).

Therefore, we sought to examine this pathway for processing innate threats as well, with the view that a side-by-side comparison may lead to a new mechanistic insight into the similarities and differences between circuits processing innate and learned threats.

Here, we show that, as with the learned threat conditioning, the MGN-BLA pathway is essential for processing the innately aversive looming stimulus. Inactivation of either the BLA or the BLA projecting neurons in the MGN was sufficient to impair defensive responses to innate and learned threats. Additionally, fiber photometry from the BLA neurons or the MGN axons projecting to the BLA showed a rapid rise in their activity to both forms of stimuli. However, the activity was reduced as mice showed habituation to the aversive stimuli. Despite similarities in processing the innate and learned threat, we found that propranolol, an beta-adrenergic receptor blocker, specifically impairs the innate defensive response, while the response to the learned threat remains intact.

## Results

### BLA activity is required for processing a visual innately aversive threat and aversive conditioning

Threat conditioning is an associative learning paradigm where an initially neutral tone (conditioned stimulus, CS) is repeatedly paired with an aversive foot-shock (unconditioned stimulus, US). The next day, mice, upon exposure to the CS, show freezing responses (conditioned response, CR), indicating successful learning of the association (Blair et al., 2001; Pape & Pare, 2010).

As for the innately aversive threat, we used the looming stimulus, an overhead expanding black disk that is thought to mimic an approaching aerial predator (Yilmaz & Meister, 2013). Unlike the tone used in threat conditioning, the looming stimulus triggers defensive responses without prior learning. The defensive response may vary from freezing to escapes and tail rattling (Yilmaz & Meister, 2013; Salay et al., 2018). In this study, we tested mice in an arena devoid of shelter (Shang et al., 2018). This setup promotes freezing as the dominant defensive response, comparable to the freezing response observed in threat conditioning. Although the mice showed extended freezing beyond the looming stimulus period, we observed rapid habituation to the repeated presentation of the loom (Figure S1A-C).

It is well established that the BLA activity is required to process learned threats (Maren et al., 1996; Maren, 1999; Anglada-Figueroa & Quirk, 2005; Johansen et al., 2014). To test whether the BLA is required for visually-evoked innate defensive response, we applied a loss-of-function approach by transiently inactivating the BLA. We infected the BLA pyramidal neurons with hM4Di (Gi-coupled human muscarinic M4 designer receptor exclusively activated by a designer drug (iDREADD)) tagged with m-Cherry fluorescent proteins. We validated that the efficacy of iDREADDs’ agonist, Clozapine-N-oxide (CNO), mediated inhibition by performing optical stimulation of MGN axons and in vivo electrophysiology recording in the BLA (Figure 1A). Intraperitoneal injection of CNO reduced the light-evoked BLA activity significantly (Figure 1B) (Armbruster et al., 2007; Stachniak et al., 2014). Behaviorally, the iDREADD-mediated BLA silencing significantly reduced the defensive responses of the animals to the looming stimulus (Figure 1C,D and S1D-H).

**Figure 1.**
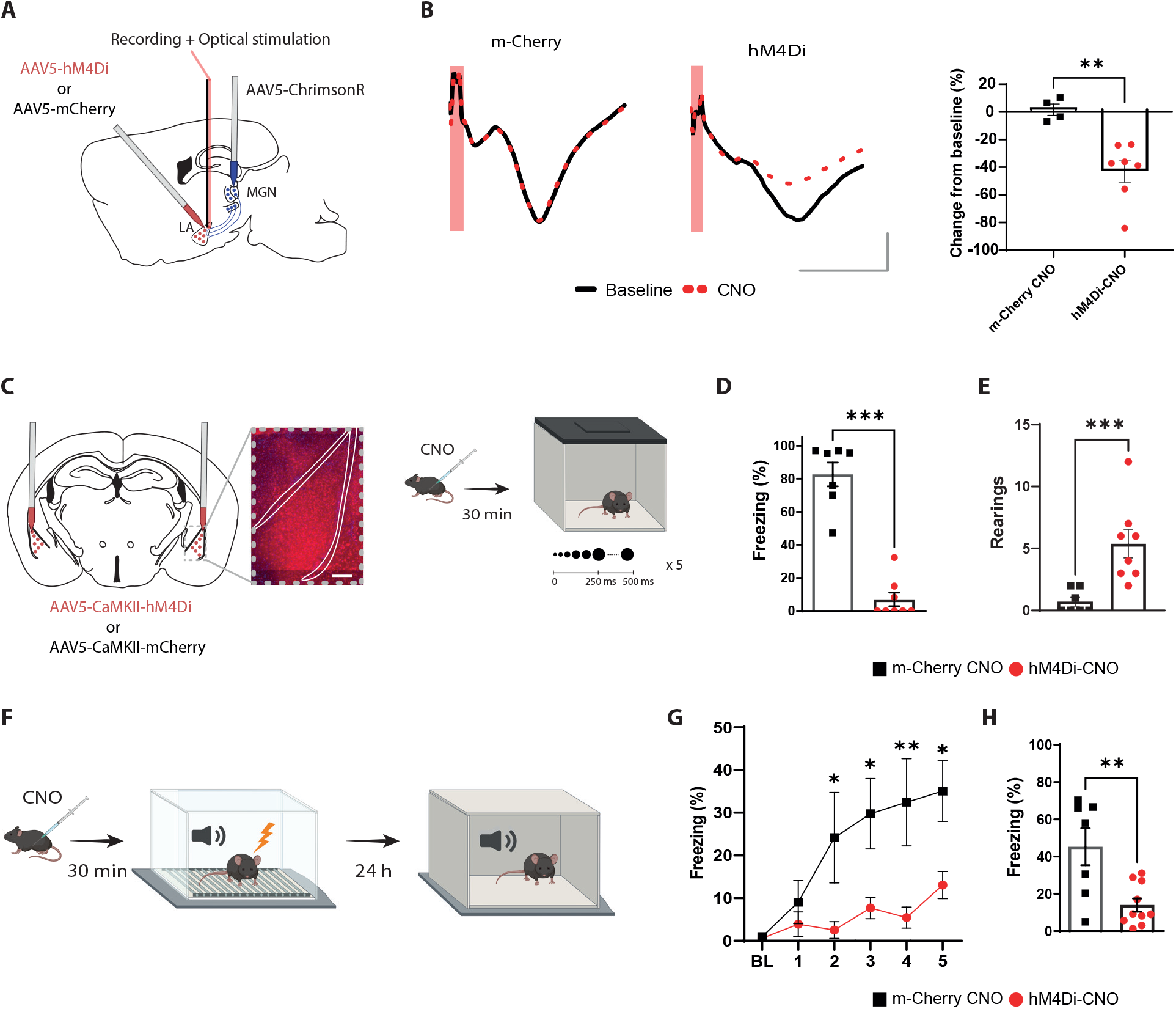
The BLA activity is required for processing innate as well as learned aversive signals. **A)** Experimental design of the in-vivo electrophysiology experiment. Mice were injected unilaterally with AAV vectors expressing ChrimsonR in the MGN and with hM4Di or m-Cherry in the BLA. **B)** CNO reduces the amplitude of the fEPSP in hM4Di-but not m-Cherry expressing neurons. Left panel: fEPSP is unchanged after CNO injection in the m-Cherry group. Middle panel: fEPSP is reduced after CNO injection in the hM4Di group. Scale bar, 5 ms, 0.1 mV, The red bar represents the pulse of light (0.5 ms, 638 nm). Right panel: Differential score comparing the change between before and after CNO injection in the two groups (m-Cherry-CNO, n=4; hM4Di-CNO, n=7; Unpaired t-test, p-value=0.0033). **C)** Experimental design of the behavioral experiment. Mice were injected bilaterally with AAV expressing hM4Di or m-Cherry in the BLA. Scale bar, 250um. After three weeks of virus expression, the mice were exposed to the looming stimulus 30 minutes after CNO injection. **D)** The freezing level is significantly reduced in the hM4Di-CNO group (n=8) compared to the m-Cherry-CNO group (n=7; Mann-Whitney test, p-value=0.0003). **E)** The rearing events are significantly reduced in the hM4Di-CNO group (n=8) compared to the m-Cherry-CNO group (n=7; Mann-Whitney test, p-value=0.0006). **F)** One day after the looming exposure, the same animals were injected with CNO 30 minutes prior to the aversive conditioning protocol. Twenty-four hours later, the mice were tested in a new context in a CNO-free trial. **G)** Freezing level during the baseline period (BL) and the five tone and foot-shock pairings. The CS-evoked freezing is significantly reduced in hM4Di-CNO group (n=10) than the m-Cherry-CNO group (n=7; Repeated-measures ANOVA for group by time interactions, F: 5,80=3.916, p-value= 0.0032 with Sìdak test correction). **H)** The CS-evoked freezing in a new context is significantly reduced in the hM4Di-CNO group (n=10) compared to the m-Cherry-CNO group (n=7; Unpaired t-test, p-value=0.0041). Results are reported as mean ± S.E.M. *, p<0.05; **, p<0.01; ***, p<0.001.

Specifically, BLA inhibition resulted in failure of switching from rearing behavior to freezing, which is a typical looming stimulus evoked behavior (Figure 1E and S1D,E). The following day, the same mice were subjected to the threat conditioning protocol in the presence of CNO (Figure 1F). Consistent with previous reports (Angla-da-Figueroa & Quirk, 2005; Johansen et al., 2014; Maren, 1999; Maren et al., 1996), transient inactivation of the BLA during the conditioning reduced the freezing response to the CS during the conditioning as well as the recall periods (Figure 1 G,H and S1I,J). Neither the injection of CNO nor the expression of iDREADDs alone interfered with the expression of the defensive responses to innately aversive and learned threats. In addition, mice with a permanent lesion in the BLA showed a similar deficit in defensive responses (Figure S2). Thus, the neuronal activity within the BLA is not only required for encoding and processing the learned threat but it is also essential for processing an innately aversive threat. This, to our knowledge, is the first direct evidence documenting that the BLA is required for processing an innate visual threat cue.

### The selective lesion of the BLA-projecting MGN neurons impairs the defensive responses to the looming stimulus and to the aversive conditioning

The subcortical pathway comprising the MGN inputs to the BLA is known to be essential for the acquisition of threat conditioning (Romanski and LeDoux, 1992 a and b; LeDoux et al., 1984; Campeau & Davis, 1995; Barsy et al., 2020; Taylor et al., 2021; Lee et al., 2021). Therefore, we tested whether these inputs are also required for processing innately aversive threats. We selectively lesioned BLA-projecting MGN neurons by injecting retroAAV2-Cre in the BLA and a mixture of DIO-GFP and DIO-taCaspase3 in the MGN (Yang et al., 2013) (Figure 2A). Additionally, in two separate control groups, we injected retroAAV2-Cre in the BLA and DIO-GFP in the MGN (GFP group) or a mixture of DIO-GFP and DIO-taCaspase in the MGN (sham group). Our retrograde labeling was largely confined within the MGN (Figure 2B,D).

**Figure 2.**
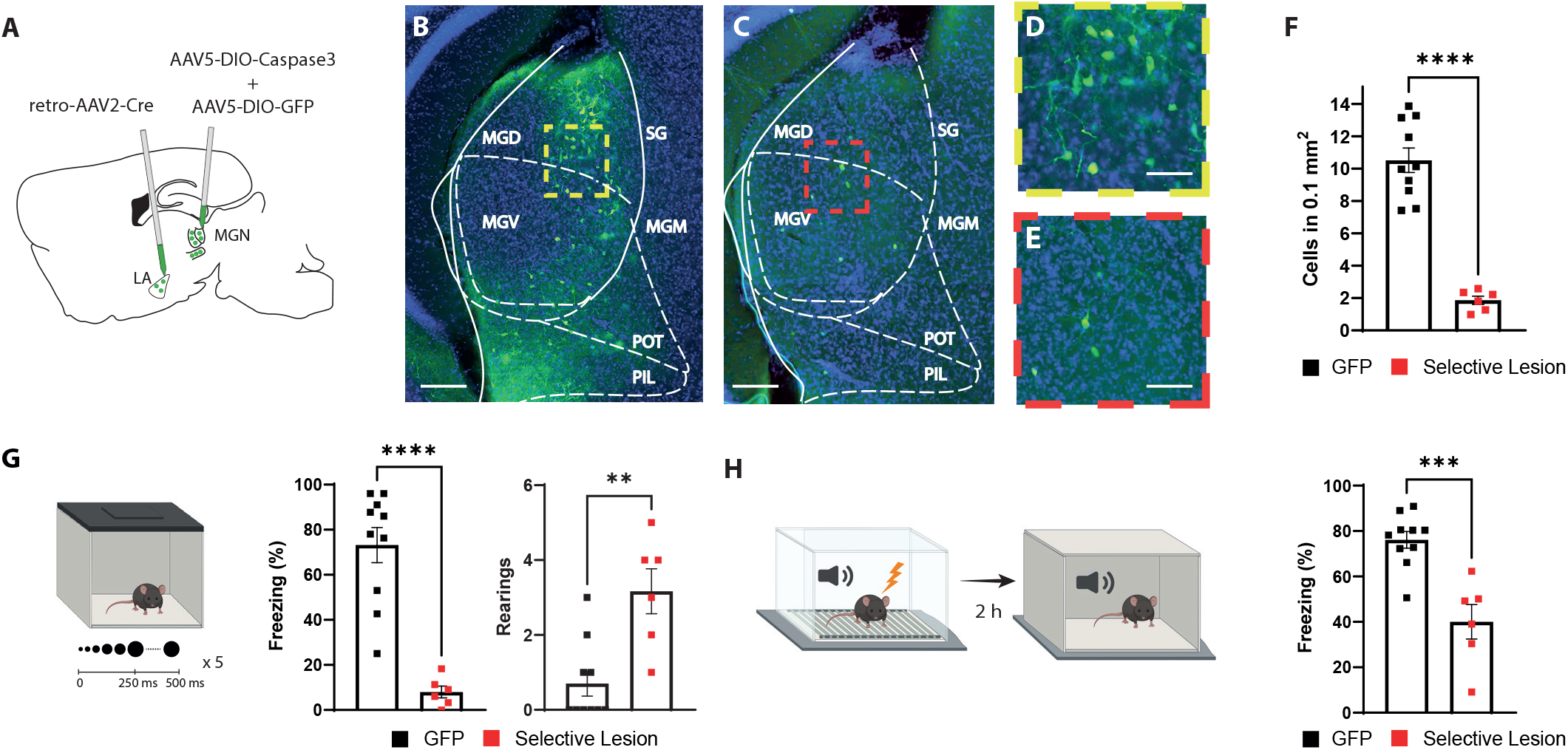
Selective lesion of the BLA-projecting neurons in the MGN impairs the defensive responses to the looming stimulus and to the aversive conditioning. **A)** Viral strategy for the selective lesion of BLA-projecting MGN neurons. The GFP group was injected with retro-AAV2-Cre in BLA and AAV5-DIO-GFP only in the MGN. **B,C)** Representative image showing retrogradely transported GFP in the BLA-projecting MGN neurons in a mouse from the GFP group **(B)** and a mouse from the Selective lesion group **(C)**, respectively. Scale bar, 200um. **D,E)** Zoomed-in images from the regions outlined in yellow from **B** and in red from **C**. Scale bar, 50um. **F)** Quantification of GFP+ neurons in the MGN. The selective lesion group (n=6) showed a significant reduction in the number of GFP+ neurons compared to the GFP group (n=10; Unpaired t-test, p-value<0.0001). **G)** Mice were exposed to the looming stimulus. The freezing level is significantly reduced in the selective lesion group (n=6) compared to the GFP group (n=10; Mann-Whitney test, p-value<0.0001). The rearing frequency is significantly higher in the selective lesion group (n=6) compared to the GFP group (n=10; Mann-Whitney test, p-value=0.0050) **H)** The same mice were conditioned one day after the looming exposure. The mice were tested for memory recall in a new context two hours later. The selective lesion group (n=6) showed a significant reduction in CS-evoked freezing compared to the GFP group during the STM recall (n=10; Unpaired t-test, p-value=0.0003). Results are reported as mean ±S.E.M. **, p<0.01; ***, p<0.001; ****, p<0.0001.

Among mice with the selective lesion of the BLA-projecting MGN neurons, we observed few GFP+ neurons (Figure 2C,E,F), demonstrating the efficiency of the approach. Furthermore, mice with the selective lesion showed a significant reduction in their defensive response to the looming stimulus as opposed to the two control groups (Figure 2G and S3A-C). Similar to the BLA inhibition experiment, the selective lesion caused a similar failure in switching from exploratory to defensive behavior upon looming stimulus exposure (Figure 2G and S3A,B). Likewise, the defensive responses to the learned aversive cue were reduced during the conditioning as well as the recall sessions (Figure 2H and S3D,E). Together these data, in line with a recent report (Kang et al., 2022), demonstrate that the activity of the BLA-projecting MGN neurons is required for the defensive responses not only to a learned, but also to an innately aversive threat.

### The axons of the BLA-projecting MGN neurons are activated by the looming stimulus and show an increase in CS-evoked response following aversive conditioning

In the preceding section, we demonstrated that the activity of the BLA-projecting MGN neurons is essential for processing innate and learned threats. Therefore, we expect an increase in the activity of the MGN input to the BLA that is time-locked to the threat signals. For this purpose, we took advantage of fiber photometry in freely moving mice. Virus expressing the genetically encoded Ca^2+^ indicator GCaMP7s (Dana et al., 2019) was injected into the MGN, and a fiber optic was implanted above the dorsal tip of the BLA (Figure 3A-C). The axonal activity of the MGN neurons serves as a proxy for their release of neurotransmitters into the BLA. The time-locked GCaMP activity of the MGN projections to the onset of the looming stimulus was evident. As mice showed habituation to the stimuli, GCaMP activity diminished, with later stimuli eliciting neither defensive behavior nor time-locked GCaMP activity in the MGN inputs (Figure 3D-J).

**Figure 3.**
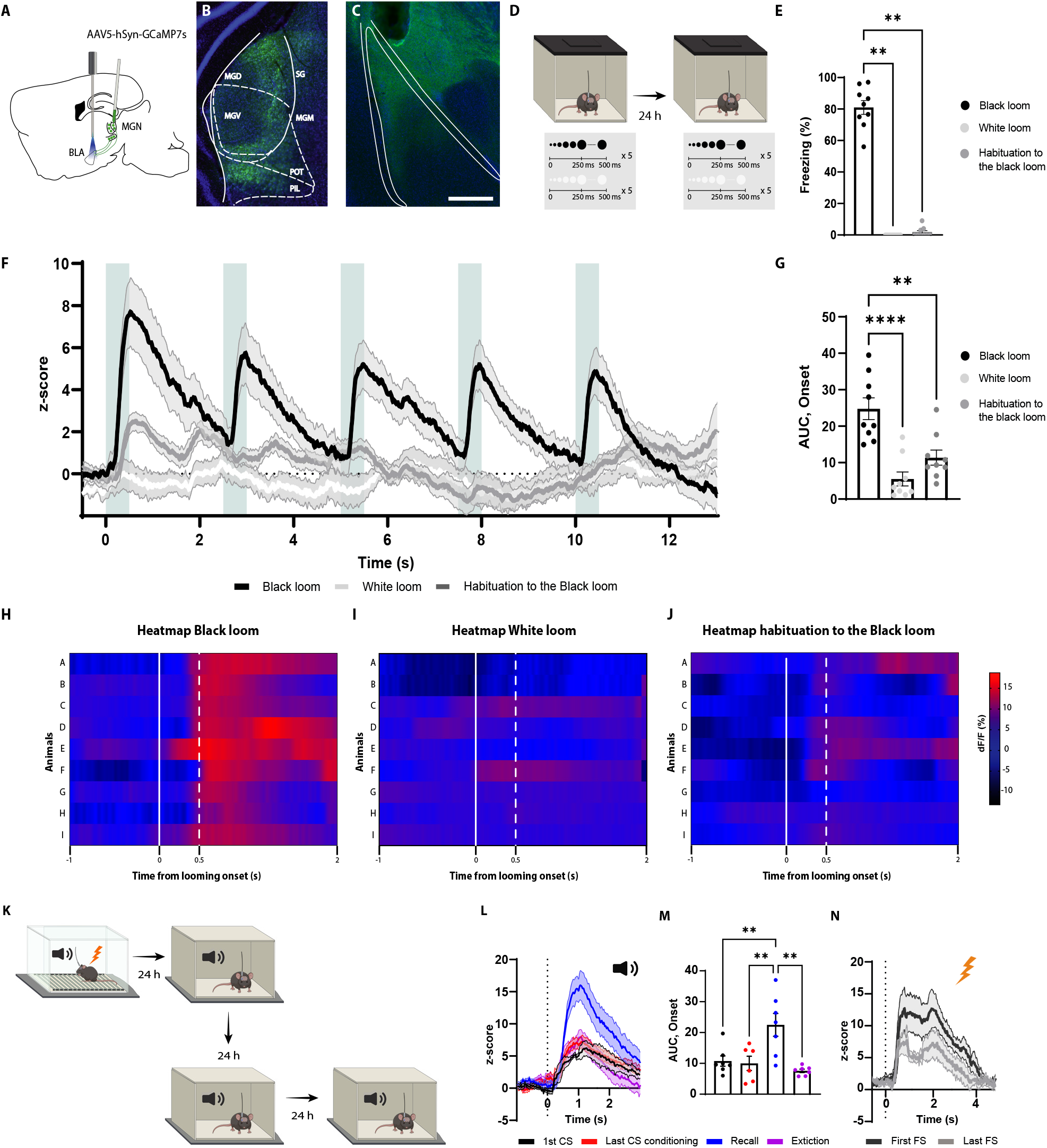
The axons of the BLA-projecting MGN neurons are activated by the looming stimulus and show an increase in CS-evoked response following aversive conditioning. **A)** Illustration showing virus injection and the optic fiber implantation strategy for GCaMP7s recordings from the axon terminals of the BLA-projecting MGN neurons. **B)** Representative image of GCaMP7s expression in the MGN. Scale bar, 200um. **C)** Representative image of the optic fiber location above the BLA. Scale bar, 500um. **D)** After five weeks of virus expression, the mice were exposed to the black and white looming stimulus. Twenty-four hours later, the mice were reexposed again to the looming stimulus. **E)** Freezing level to the black looming stimulus (n=9), to the white looming stimulus (n=9) and after the habituation to the black looming (n=9, Friedman test pvalue<0.0001 with Dunn’s test).**F)** Z-score of the Ca2+ response from the MGN axon terminals during the black looming stimulus (n=9; in black), during the white looming stimulus (n=9; in white) and during the habituation to the black looming stimulus (n=9; in grey). **G)** The area under the curve (AUC) is significantly reduced when the mice are habituated to the black looming compared to the first exposure to it (n=9; Paired t-test, p-value=0.0001). **H-J)** Heatmap of the response to the first expansion of the black looming stimulus (**H**), white looming stimulus (**I**), and after the habituation to the black looming stimulus (**J**) for each mouse. **K)** Mice were conditioned and tested as previously described. After the recall session, the same went through an extinction protocol for the following two days. **L)** Z-score of the Ca2+ response from the axon terminals of the BLA-projecting MGN neurons during the first CS presentation (n=7; in black), during the last CS presentation of the conditioning (n=6; in red), during the first CS presentation during the recall session (n=7; in blue) and during the first CS presentation after the extinction training (n=7; in purple). **M)** The AUC is significantly increased when the mice are exposed to the CS during the recall session (n=7) compared to the CS-evoked response at the beginning of the conditioning (n=7) and at the end of the conditioning (n=7) and after the extinction training (n=7; Mixed-effects analysis, F: 3,17 = 8.791, p-value= 0.0010 with Tukey test correction). **N)** Z-score of the Ca2+ response from the axon terminals of the BLA-projecting neurons in the MGN during the first footshock presentation (n=7). Results are reported as mean ±S.E.M. **, p<0.01; ***, p<0.001.

Notably, the white looming stimulus, as opposed to a black looming stimulus, did not evoke defensive behavior (Yilmaz & Meister, 2013), nor did it trigger GCaMP activity (Figure 3E-I). This, along with the result from the habituation sessions, indicates that the increased activity of the MGN inputs is not merely the product of the sensory property of the looming stimulus, but it reflects the saliency and aversiveness of the stimulus, as well.

We next monitored the activity of the MGN inputs during aversive conditioning, the recall sessions, and post-extinction training (Figure 3K). As expected from a multisensory brain region (Bordi & LeDoux, 1994; Linke et al., 1999; Linke, 1999), the MGN inputs were activated from the tone onset (Figure 3L,M). The amplitude of the activity remained unchanged for the subsequent CS presentations, despite mice showing CS-evoked freezing to these stimuli (Figure 3L). Interestingly, during the recall session 24 hours later, we observed a significant increase in CS-evoked GCaMP activity, which was not evident during the conditioning (Figure 3L,M). In addition, upon extinction training, the CS-evoked activity returned to its preconditioning level (Figure 3L,M). Moreover, the footshock induced a time-locked increase in GCaMP activity, which was reduced in amplitude with subsequent US delivery (Figure 3N). From these experiments, we conclude that the MGN neurons directly convey the signals for innately aversive as well as for learned threats to the BLA.

### Contralateral disconnection of the MGN-BLA pathway impairs the defensive responses to the looming stimulus and to the aversive conditioning

Our inactivation experiments demonstrate that the BLA (Figure 1) as well as the BLA-projecting neurons in the MGN (Figure 2) are necessary for the processing of the learned and innate aversive threat responses. However, the previous experiments on their own cannot distinguish whether the two regions function in series, with the BLA receiving the threat signals from the MGN (as indicated by GCaMP activity in the MGN inputs) (Figure 3); or, the MGN and the BLA function in parallel, with the BLA receiving the signal from other sources. To address this issue, we used an asymmetrical disconnection approach (LeDoux et al., 1986; Iwata et al., 1986; Eldridge et al., 2015; Barker et al.; 2017; Torromino et al., 2019), where we inhibited the activity of the MGN and the BLA contralaterally. This approach is suited to test whether the direct connection between two regions is required for a particular function (Figure 4A,B). The prerequisite is the connections should be ipsilateral and not reciprocal, as it is the case for the MGN and the BLA (LeDoux et al., 1986; LeDoux, 1990).

**Figure 4.**
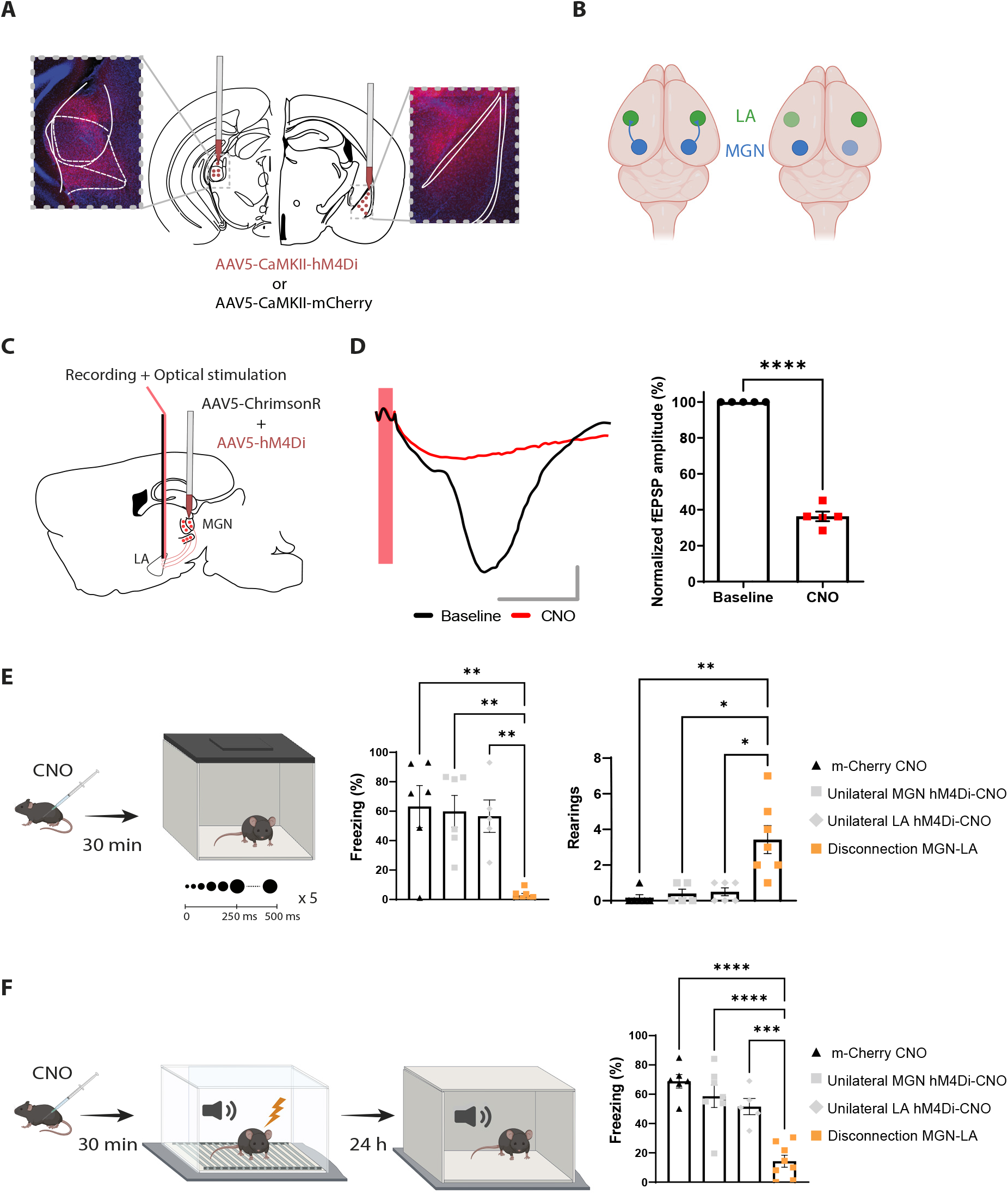
Reversible contralateral disconnection of the MGN-BLA pathway impairs the defensive responses to the looming stimulus and to the aversive conditioning. **A)** hM4Di injections in the MGN and the contralateral BLA. **B)** Left: the MGN and the BLA are connected through a non-reciprocal and ipsilateral connection. Right: contralateral disconnection of the MGN-BLA pathway. **C)** Diagram showing the experimental design of the in vivo electrophysiology experiment. Mice were co-injected unilaterally with AAV vectors expressing ChrimsonR and hM4Di in the MGN. **D)** CNO reduced the fEPSP in mice expressing hM4Di and injected with CNO. Left panel: Representative traces from one mouse from the hM4Di-CNO group. Scale bar, 5 ms, 0.1 mV. Right panel: the graph shows the normalized fEPSP values before and after CNO injection in mice expressing hM4Di and injected with CNO (n=5; Paired t-test, p-value<0.0001). **E)** Mice were injected with CNO thirty minutes before being exposed to the looming stimulus. The Disconnection MGN-BLA group (n=7) showed a significant reduction in the freezing levels compared to all the groups. The unilateral inhibition of the MGN (n=6) and the BLA (n=5) did not impair the looming stimulus-evoked freezing (Ordinary one-way ANOVA, F=3,20, p-value=0.0006). Right: The rearing events are significantly higher in the Disconnection MGN-BLA group (n=7) compared to all the other groups during the looming stimulus presentation (Kruskal-Wallis test, F=2,24, p-value=0.0016). **F)** Mice were reinjected with CNO thirty minutes before aversive conditioning, and they were exposed to the CS in a new context in a CNO-free trial. The Disconnection MGN-BLA group (n=7) showed a significant reduction in the CS-evoked freezing level compared to all control groups during the LTM recall (Ordinary one-way ANOVA, F=3,22, p-value<0.0001).Results are reported as mean ±S.E.M. *, p<0.05; **, p<0.01; ***, p<0.001; ****, p<0.0001.

For this purpose, we applied a reversible disconnection between the MGN-BLA pathway by expressing hM4Di in the MGN and the BLA contralateral to each other (Figure 4A,B). Electrophysiologically, upon CNO injection, the responses of the BLA neurons to the optical stimulation of MGN neurons co-expressing ChrimsonR and hM-4Di was greatly reduced (Figure 4C,D). Behaviorally, CNO-induced inactivation of the contralateral regions significantly reduced the de-fensive responses to the looming stimulus (Figure 4E and S4A-H) and to the recall session of the aversive conditioning (Figure 4F and S4I-K). CNO injection in mice expressing mCherry in the contralateral MGN and BLA did not reduce freezing responses in either of the behavioral tasks (Figure 4E,F). More importantly, unilateral inactivation of the MGN and the BLA was not sufficient to block the defensive responses (Figure 4E,F). Similar results were obtained when we performed an irreversible disconnection of the MGN-BLA pathway (Figure S4L-O). These experiments suggests that the direct projection from the MGN to the BLA is required for the processing of innately aversive threats as well as learned threats.

### The MGN lesion blocks the BLA neuronal activation by the looming and conditioned stimuli as well as it reduces the response to a foot shock

The disconnection of the MGN-BLA pathway impairs the defensive responses to both forms of threat cues (Figure 4), suggesting that this pathway is the main root by which the BLA receives the aversive signals. If so, upon the MGN lesion, the BLA responses to the aversive stimuli must largely disappear. We co-injected AAV vectors expressing DIO-taCapsase3 and Cre recombinase in the MGN and GCaMP8m in the BLA (Zhang et al., 2021). GCaMP signal was collected through a fiber optic implanted above the tip of the BLA (Figure 5A and S5A). The control group underwent the same procedure except that no Cre recombinase was injected.

**Figure 5.**
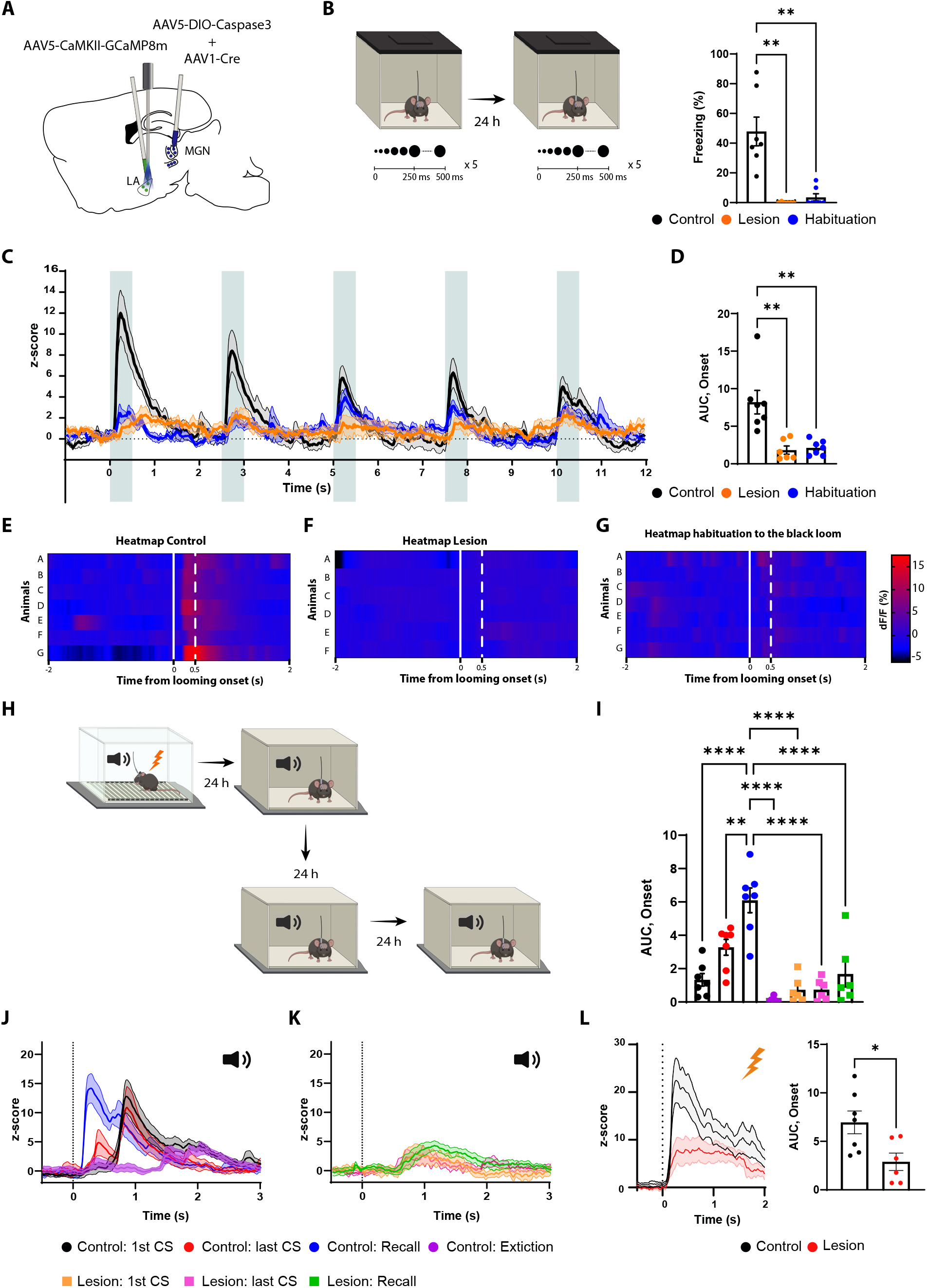
The MGN lesion impairs the BLA response to the looming, the CS, and the US stimuli. **A)** Illustration showing the virus injections and the optic fiber implantation. **B)** Mice were exposed to the looming stimulus on day 1. On day two the control group was habituated to the looming stimulus. The mice from the lesion group (n=6) have a significant reduction in the freezing levels to the looming stimulus compared to the Control group (n=7). Similarly to the control group, once the mice were habituated to the black looming stimulus (n=7; Kruskal-Wallis test, F= 3,20, p-value<0.0001). **C)** Z-score of the calcium response during the black looming stimulus in the control group (n=7; in black) and in the Lesion group (n=6; in orange) and the control group after the habituation to the black loom (n=7; in blue). **D)** The AUC is significantly reduced in the lesion (n=6) and in the habituation to the black loom (n=7) groups compared to the control group (n=7; Ordinary one-way ANOVA, F (2,17) = 12.73, p-value=0.0004) during the exposure to the black looming stimulus. **E-G)** Heatmap representing the individual response to the first expansion to the black looming stimulus for the control group (**E**) and for the lesion group (**F**), and for the control group after habituaion to the black loom (**G**) for each mouse. **H)** Mice were conditioned and tested as previously described. After the recall session, the same went through an extinction protocol for the following two days. **I)** The AUC for the first CS presentation during the recall in the control group is significantly increased compared to all conditions of the lesion group (Two-way ANOVA, F: 6,33=16.29, p-value< 0.0001 with Tukey test correction). **J)** Z-score of the calcium responses during the first (n=7) and last CS (n=7) presentation during the conditioning (in black and in red), and during the first CS presentation during the recall (n=7; in blue) and after extinction training (n=7; in purple) in the control group. **K)** Z-score of the calcium responses during the first (n=6) and last CS (n=6) presentation during the conditioning (in orange and in magenta), and during the first CS presentation during the recall (n=6; in green) in the lesion group. **L)** Left: Z-score of the Ca2+ response during the foot shock presentation for the control group (n=7, in black) and the Lesion group (n=6, in red). Right: The AUC is significantly reduced in the mice from the lesion group (n=6) compared to the mice from the control group (n=7; Unpaired t-test, p-value=0.0219). Results are reported as mean ±S.E.M. *, p<0.05; **, p<0.01; ****, p<0.0001.

In the MGN-lesioned mice, the BLA response to the black looming stimuli, along with the behavioral defensive responses, largely disappeared (Figure 5B-F). In the non-lesioned mice, where defensive responses to the looming stimuli remained intact, we observed a timed-locked GCaMP activity to the stimuli in the BLA (Figure 5B-E). After habituation, the behavioral and neuronal responses to the black looming stimulus in the control group were comparable to those lesioned (Figure 5B-G). Moreover, the white looming stimulus did not produce a noticeable activation of the BLA neurons, nor did it elicit a defensive response (Figure S5B-D).

We next monitored the activity of the BLA during the auditory threat conditioning and the recall in the MGN-lesioned and non-lesioned mice (Figure 5H). The MGN-lesioned mice, as expected, did not produce a conditioning response to the tone (Figure S5F,G). Accordingly, the CS failed to activate the BLA in these mice (Figure 5I-K and S5E), while the US-evoked response was significantly reduced (Figure 5L). In the non-lesioned mice, on the other hand, we observed tone-evoked response in the BLA, but with a latency that cannot be fully explained by the slow kinetics of the calcium indicator. As the conditioning progressed, we observed the appearance of an additional, smaller but shorter latency tone-evoked component. In the recall session, the tone-evoked response in the BLA was significantly larger in amplitude, and shorter in latency than observed at the end of the conditioning session in the previous day (Figure 5J). Upon extinction, the tone-evoked response, along with the defensive behavior, was greatly reduced (Figure 5I,J and S5G).

### Blocking beta-adrenergic receptors reduce the defensive response to the innately aversive threat

Previous studies have shown that innately aversive stimuli such as fox urine (Hu et al., 2007; Liu et al., 2010) or cat fur odor (Do Monte et al., 2008) mediates defensive responses through the activation of the beta-adrenergic receptor. Therefore, we considered that the threat response triggered by looming stimulus may rely on the activity of these receptors. To test this, prior to exposure to the looming stimulus mice were injected with propranolol, a beta-adrenergic receptor blocker (Figure S6A-C). The defensive responses to the looming stimulus in these mice largely disappeared, while the exploratory behavior remained intact (Figure S6B). On the other hand, Sotalol, a peripherally acting beta-adrenergic blocker, had no impact on defensive responses, suggesting propranolol reduced the looming stimulus-evoked defensive responses by acting on the central nervous system (Figure S6B).

To further examine the contribution of the subcortical pathway in the processing of innate threats, we injected propranolol in mice expressing GCaMP in either MGN axons projecting to the BLA or in the BLA pyramidal neurons, followed by repeated exposure to looming stimuli (Figure 6A). At the cellular level, looming stimuli failed to elicit a significant increase in the BLA activity, which reflects the reduced freezing response in mice injected with propranolol (Figure 6B-D). Surprisingly, the stimulus-evoked MGN axonal activity remained undisturbed despite the lack of defensive response (Figure 6E-G). Of note, 48 hours later, when these mice were re-tested, defensive response, as well as time-locked BLA activity to the looming stimulus, was restored (Figure 6B-E).

**Figure 6.**
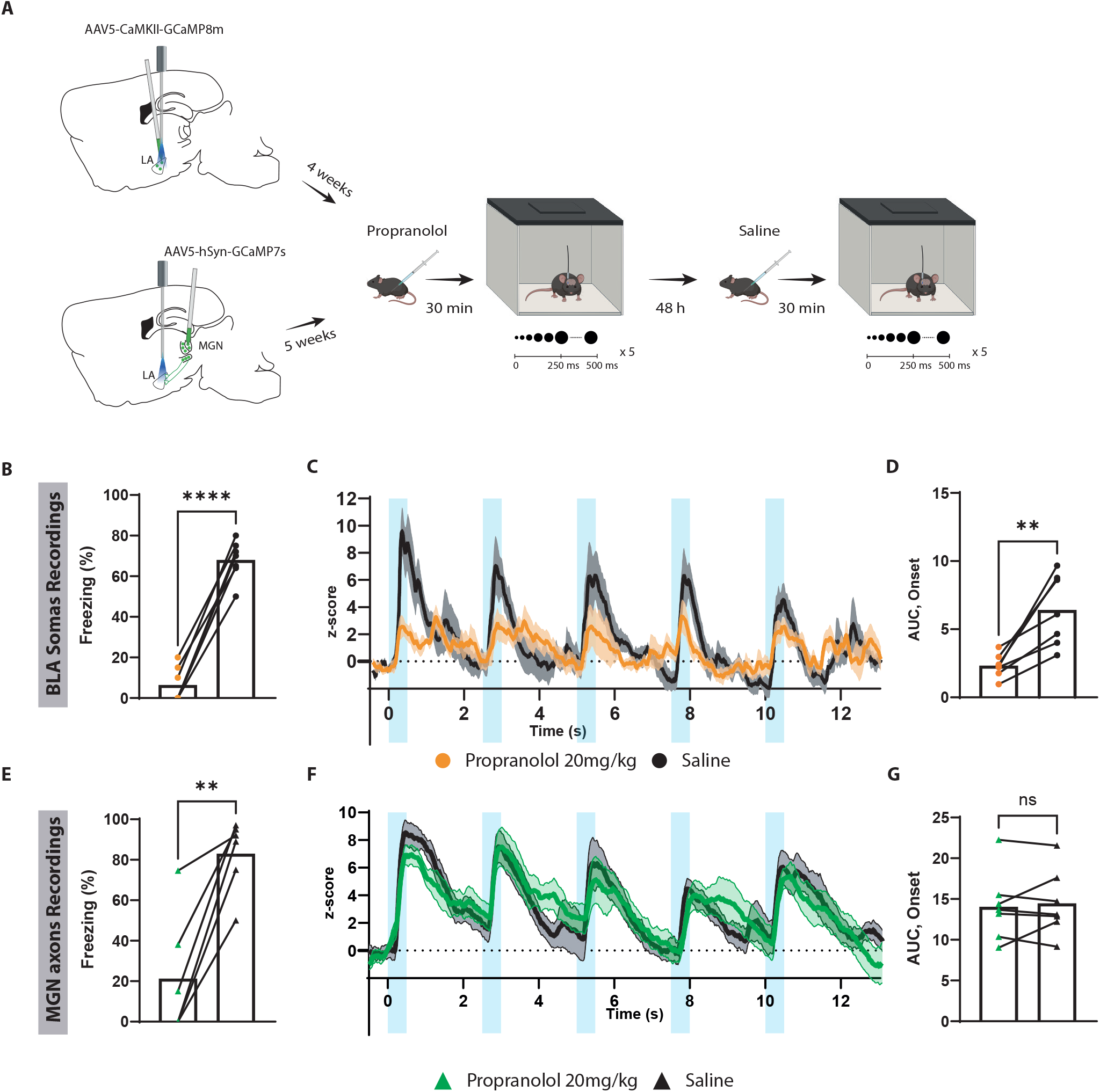
Blocking beta-adrenergic receptors reduce the defensive response and the BLA activation to the innately aversive threat. **A)** Timeline showing the different stages of the experiment. **B)** The looming stimulus-evoked freezing is significantly reduced in the propranolol trial compared to the re-exposure trial in which the same mice were injected with saline (n=7; Paired t-test, p-value<.0001). **C)** The graph shows the average of the z-score of the Ca2+ responses of the looming stimulus presentation in the BLA after propranolol injection (n=7, in orange) and saline injection (n=7; in black). **D)** The AUC is significantly reduced in the propranolol trial (n=7) compared to the saline trial (n=7; Paired t-test, p-value=0.0034). **E)** The looming stimulus-evoked freezing is significantly reduced in the propranolol trial compared to the re-exposure trial in which the same mice were injected with saline (n=7; Paired t-test, p-value=0.0022). **F)** The graph shows the average of the z-score of the Ca2+ responses of the looming stimulus presentation of the MGN axon terminals after propranolol injection (n=7, in green) and saline injection (n=7; in black). **G)** The AUC is unchanged in the propranolol trial (n=7) compared to the saline trial (n=7; Paired t-test, p-value=0.6016). Results are reported as mean ±S.E.M. ns, non-significant; **, p<0.01; ****, p<0.0001.

To our surprise, propranolol did not impair the aversive conditioning responses when injected either before the conditioning or before the recall session (Figure S10D-I). In addition, propranolol injections prior to the recall session perturbed neither BLA (Figure S6J-M) nor MGN axonal activity (Figure S6N-P).

## Discussion

The thalamic-BLA pathway and its intricate microcircuitry have been described for their function in processing the auditory CS and aversive US (Barsy et al., 2020; Janak & Tye, 2015; Rogan et al., 1997; Rogan et al., 2005). Yet, their role in processing unimodal innate threats remains underinvestigated (Kang et al., 2022). Here, we demonstrate that transient or permanent inactivation of the BLA (Figure 1) or BLA-projecting MGN neurons (Figure 2) not only impairs threat learning but also it abolishes all the measured defensive responses to an innate threat. More specifically, upon exposure to an innate visual threat, animals with a compromised MGN-BLA pathway fail to switch from exploratory to defensive behavior (Figure 1,2,4). Furthermore, the MGN axons in the BLA (Figure 3F) and the neurons within the BLA produce time-locked activity to each looming stimulus in mice showing defensive responses (Figure 5C). With repeated exposure to looming stimuli, the neuronal activity, along with defensive responses, gradually fades.

An overlap in processing innate and learned threats is not unique to the MGN-BLA pathway. The lateral habenula and the central amygdala control the processing of a number of innate and learned threats (Fadok et al., 2017; Lecca et al., 2017; Lecca et al., 2020; Matsumoto & Hikosaka, 2007; Root et al., 2014; Mondoloni et al., 2022; Tovote et al., 2016; Sachella et al., 2022; Isosaka et al., 2015). Since both types of threats share a similar repertoire of defensive responses, such as freezing and escaping, the wiring economy favors the layout where there is closer physical proximity between the two circuits, as we observed here (Klyachko and Stevens, 2003; Stevens, 2012).

Direct and indirect evidence show that the MGN-BLA pathway processes other forms of innate threats as well. Recently, studies have demonstrated that inactivation of thalamic-BLA pathway reduces freezing response to intense sound and looming stimuli (Kang et al., 2022), as well as innately aversive ultrasound activates the BLA (Mongeau et al., 2003; Shukla and Chattarji, 2022). Moreover, the use of live predators as an innate threat has further supported this notion by showing that the BLA lesion in rodents eliminates defensive responses to the threat (Bindi et al., 2018; Martinez et al., 2011).

Since the MGN is widely regarded as an auditory relay region (Rogan & LeDoux, 1995; Rogan et al., 1997; Weinberger, 2011), its role as the main source of innately aversive visual signal to the BLA may seem unexpected. However, it has been known that the MGN receives auditory, somatosensory, visual, and multimodal information (Bordi et al., 1994; Linke et al., 1999; Linke, 1999). The lateral posterior nucleus of the thalamus (LP) has been proposed as another direct source conveying a looming stimulus signal to the BLA (Wei et al., 2015). Our reversible disconnection of the MGN-BLA pathway, which largely spares the LP, argues otherwise. The MGN lesion not only abolishes the defensive responses to the looming stimulus but also largely eliminates the stimulus-evoked activity in the BLA (Figure 5).

Although the input source of the looming stimulus to the MGN is unknown, we speculate the superior colliculus (SC) could be a likely candidate as it encodes threat and escape behavior to looming stimuli (Evans et al., 2018) and sends monosynaptic connections to the MGN (Linke et al., 1999). Additionally, it has been shown that the SC may trigger defensive responses to the looming stimulus by conveying the signals to the periaqueductal grey (PAG) (Evans et al., 2018) or indirectly to the central nucleus of the amygdala (CeA) (Shang et al., 2015; Zhou et al., 2019). It is relevant to note that the BLA conveys learned aversive signals directly to the CeA (Fadok et al., 2017), and indirectly, through the CeA to the PAG (Tovote et al., 2016). It will be of particular interest to test whether the information signaling the innate threat is communicated through the same channel. As the SC and the BLA, the two essential regions in processing the innate visual threat, share similar downstream targets, their specific contribution to the process warrants further inquiry.

The activities of the MGN and the BLA largely mirrored each other, however, there are noticeable differences. In mice injected with propranolol, while the defensive responses to the looming stimulus were diminished significantly (Figure 6B,E), the stimulus-induced activity was reduced only in the BLA (Figure 6C), with the response of the axons of the BLA-projecting MGN neurons remaining unchanged (Figure 6F). In addition, the two regions displayed different activity in processing the learned threat. As animals learned the CS-US association, we observed an enhanced short-onset auditory response in the BLA (Figure 5J) (Quirk et al., 1995). We did not detect a similar conditioning-correlated change in the activity of the MGN axons (Figure 3L). It must be noted that because of our use of calcium indicators, we cannot exclude milliseconds change in the auditory response latency in the MGN axons. However, previous works using sub-millisecond single-unit recording also showed similar patterns (Barsy et al., 2020; Bordi & LeDoux, 1992).

Although not the focus of this work, we observed several intriguing physiological features during the conditioning and recall sessions. The conditioning increases the tone-induced activity in the BLA and reduces the response time onset (Barsy et al., 2020; Bordi et al., 1993). Remarkably, the increased response and decreased time onset were significantly more pronounced on the recall day. This cannot be attributed to the occlusion of increased firing rate caused by foot shocks, as there was no difference in the baseline activity between pre- and post-conditioning periods (data not shown). This is in line with previous studies using single-cell imaging in the BLA (Grewe et al., 2017). Regarding the increased activity, we observed a similar pattern in the MGN input where there was a significant enhanced activity during the recall session, which was not visible during the conditioning.

Our use of the beta blocker in this study was based on previous studies showing that propranolol decreases the defensive behavior to cat fur (Do Monte et al., 2008) and fox urine (Hu et al., 2007; Liu et al., 2010). We reasoned that looming stimulus, unlike a neutral tone, may elicit a defensive response in part through the activation of beta-adrenergic receptors. Our fiber photometry recording is consistent with this argument. Mechanistically, the looming stimulus may act similarly to predator odors by enhancing NE release in the brain (Hayley et al., 2001), and increasing neuronal excitability (Liu et al., 2010).

The MGN-BLA pathway typically has been evaluated in relation to associative learnings (Janak and Tye, 2015; Tye et al., 2008), especially associative learned threats (Barsy et al., 2020; Rogan et al., 1997; Rogan et al., 2005). The evidence provided here, along with a recent study (Kang et al., 2022), raises the possibility for an interaction, even a synergy, between the circuits processing innate and learned threats. Further work in this direction is needed to explore such an intriguing possibility.

## Methods

### Animals

All the procedures were performed on C57BL/6JRJ wildtype (Janvier, France). Mice were naïve and acclimated to the vivarium for at least a week before the beginning of the experiment. The mice were six to eight weeks old at the beginning of the experimental procedures. Animals were group-housed (3-4 per cage) with enriched conditions in a 12h light/dark cycle (the light switches on at 6 A.M.) with constant levels of humidity and temperature (22±1). Food and water were provided ad libitum. Behavioral experiments were conducted between 11 A.M. and 10 P.M. at Aarhus University at the Biomedicine department, Ole Worms Allé 8, Aarhus 8000. All the experimental procedures were conducted according to the Danish Animal Experiment Inspectorate.

### Stereotaxic surgery and virus expression

Mice were anesthetized using Isoflurane (IsoFlo vet 100%, Zoetis), and standard surgical procedures were used to expose the skull. For most of the experiments, stereoscope lights were not used during the surgical procedures because they reduced the behavioral responses (Figure S7). Before the surgery, the mice were injected subcutaneously with Buprenorphine 0.3 mg/mL (Temgesic, 0.1 mg/kg).

Electrolytic-induced lesion of the BLA: mice were lesioned bilaterally in the BLA at the following coordinates: anteroposterior (AP): −1.4 mm, mediolateral (ML): ±3.6 mm, dorsoventral (DV): −3.85/-4.1/4.35 mm from the skull. The electrolytic lesion was performed with a concentric bipolar electrode (50691, Stoelting, USA) by delivering a constant direct current at each location (0.6 mA for 15 seconds). In another group of mice, the electrode was placed at the same location without delivering any current (sham surgery).

Lesion of the BLA: mice were bilaterally injected with a mixture of AAV-5/2-hEF1α-dlox-(pro)taCasp3_2A_TEVp(rev)-dlox (titer: 4.7 x 10-12 vg/ml) in AAV-1/2-hCMV-chI-Cre- (titer: 1.0 x 10-13 vg/ ml, ratio 7:1.5). The volume of injection was 0.5 μL per hemisphere at the following coordinates AP: −1.6 mm, ML: ±3.45 mm, DV: −3.5/-4.1 mm from the skull. Control mice were injected with AAV-5/2-hEF1α-dlox-(pro)taCasp3_2A_TEVp(rev)-dlox at the same coordinates.

Chemogenetics inhibition of the BLA: mice were bilaterally injected with AAV-5/2-mCaMKIIα-hM4D(Gi)_mCherry (titer: 8.9 x 10-12 vg/ml) or with AAV-5/2-mCaMKIIα-mCherry (titer: 6 x 10-12 vg/ ml, diluted 1:1 in PBS). The injection volume was 1 μL per hemisphere at the following coordinates for the BLA AP: −1.6 mm, ML: ±3.45 mm, DV: −3.5/-4.1 mm from the skull.

Lesion of the MGN: mice were bilaterally injected with a mixture of AAV-5/2-hEF1α-dlox-(pro)taCasp3_2A_TEVp(rev)-dlox (titer: 4.7 x 10-12 vg/ml) in AAV-1/2-hCMV-chI-Cre- (titer: 1.0 x 10-13 vg/ ml, ratio 7:1.5). The volume of injection was 1 μL per hemisphere at the following coordinates at AP: −3.15mm, ML: ±1.85 mm, DV: −3.4/-3.5 mm from the skull. Control mice were injected with AAV-5/2-hEF1 α-dlox-(pro)taCasp3_2A_TEVp(rev)-dlox.

Selective lesion of the MGN projecting neurons to the BLA: animals were injected bilaterally in the MGN with a mixture of AAV-5/2-hEF1α-dlox-(pro)taCasp3_2A_TEVp(rev)-dlox in AAV-5/2-hSyn1-dlox-EGFP(rev)-dlox (titer: 1.1 x 10-13 vg/ml, ratio 7:2). The same mice were injected bilaterally with AAV-retro/2-hCMV-chI-Cre (titer: 4.4 x 10-12 vg/ml) in the BLA, using the same coordinates mentioned earlier. One control group was injected with AAV-5/2-hEF1α-dlox-(pro)taCasp3_2A_TEVp(rev)-dlox in AAV-5/2-hSyn1-dlox-EGFP(rev)-dlox (ratio 7:2) in the MGN. Another control group was injected with AAV-retro/2-hCMV-chI-Crein the BLA and with AAV-5/2-hSyn1-dlox-EGFP(rev)-dlox in the MGN using the same dilution mentioned above.

MGN-BLA disconnection experiment: mice were injected contralaterally in one MGN and one LA (randomized hemispheres). In both locations, a mixture of AAV-5/2-hEF1α-dlox-(pro)taCasp3_2A_TEVp(rev)-dlox in AAV-1/2-hCMV-chI-Cre (ratio 7:1.5). The injection volume was 1 μL per hemisphere at the coordinates described previously. For the reversible disconnection experiment: mice were injected contralaterally in one MGN and one BLA with AAV-5/2-mCaMKIIα-hM4D(Gi)_mCherry or with AAV-5/2-mCaMKIIα-mCherry (diluted 1:1 in PBS) in the BLA. Additional control groups were injected unilaterally with AAV-5/2-mCaMKIIα-hM4D(Gi)_mCherry in the MGN or in the BLA.

Fiber photometry experiments: mice were injected unilaterally with AAV-5/2-hSyn1-chI-jGCaMP7s (titer: 7.7 x 10-12 vg/ml) in the MGN or with AAV-5/2-mCaMKIIα-jGCaMP8m (titer: 6.5 x 10-12 vg/ml) in the BLA. The mice injected with jGCaMP8m in the BLA were injected in the MGN with AAV-5/2-hEF1α-dlox-(pro)taCasp3_2A_TEVp(rev)-dlox in AAV-1/2-hCMV-chI-Cre (ratio 7:1.5). The injection volume was 0.5 μL per hemisphere at the previously mentioned coordinates. Control mice were injected with AAV-5/2-hEF1α-dlox-(pro)taCasp3_2A_TEVp(rev)-dlox at the same dilution. Mono fiber-optic cannula (200/300 um, NA 0.37) was implanted in the BLA at the following coordinates AP: −1.6 mm, ML: +3.48 mm, DV: −3.45 mm from the skull. The fiber-optic cannulae were fixed to the skull with Superbond (SUN MEDICAL, Japan).

All the viral vectors were bought from Viral Vector Facility (VVF) of the Neuroscience Center Zurich (ZNZ).

### Behavioral procedures

#### Looming stimulus

The apparatus consisted of an open-top arena (37×40×19.5 cm) with a monitor (16 inches) placed on the top. No shelter was used (Barbano et al., 2020; Shang et al., 2018). The exposure to the looming stimulus was performed during the dark period (between 6 PM and 10 PM). Mice were placed in the center of the arena and explored freely for 8-10 minutes before exposure to the looming stimulus. The looming stimulus consisted of an expanding black disk over grey background, and it consisted of 5 repetitions, from 2° to 20° of visual angle (Yilmaz & Meister, 2013). The loom widens in 250 ms and remains at the same size for 250 ms with 2 seconds of pause between each loom. The experimenter was blind to the treatment and manually delivered the looming stimulus. The behavioral responses were recorded with a top camera (Phihong POE21U-1AF). All the animals were exposed at least 2-3 times to the looming stimulus, and only the one eliciting the greatest defensive response was analyzed. The defensive responses were analyzed automatically and manually using ANY-maze software (Stoelting, Ireland). An experimenter, blind to the treatment, analyzed each mouse’s freezing percentage, tail rattling, escape events and rearing events. Freezing was defined as a complete lack of movements, except for the respiratory movements, that lasted for at least 1s. Tail rattling was defined as an event in which the mouse moved the tail vigorously. Escape events were defined as a sharp increase in the locomotor speed three times greater than the average speed before the exposure to the looming stimulus. Rearing was defined as an event in which the mouse stand on its hindpaws.

For the fiber photometry experiments, a white-looming stimulus was used as a neutral control stimulus (Yilmaz & Meister, 2013). It was presented in a pseudorandom order alternated with the black-looming stimulus. The white-looming stimulus presented the same repetitions and speed as the black-looming stimulus.

#### Aversive conditioning

The apparatus consisted of an open-top cage (24×20×30 cm) with metal floor bars placed in a soundproof cubicle (55×60×57 cm) (Ugo Basile, Italy). Two different behavioral protocols were used in this study. In one, the mice were conditioned by using five pairings consisting of 20-s, 5 kHz, sine-wave tone (CS) co-terminating with a 2-s foot shock (US) 0.6 mA. On the other, the animals were conditioned using four pairings consisting of 25-s, 7 kHz, sine-wave tone co-terminating with 2-s foot shock 0.5 mA. After the conditioning, the animals stayed isolated for 10-15 minutes before returning to their homecage. Short-term (STM) and longterm memory (LTM) recall were assessed in a new context 2 hours or 24 hours after the conditioning, respectively. After 2 minutes of acclimation to the new context, the mice were presented to 4 or 5 CS presentations without the foot shock. The intertrial interval ranged between 35 s to 120s for both conditioning and testing sessions. The behavioral response was recorded by a top camera and/or side-view camera, and freezing was scored automatically by ANY-maze software.

For the fiber photometry experiments, the mice underwent an extinction protocol for two days. On each day, the CS was played twelve times in the recall context without the foot shock.

### Drugs

Propranolol hydrochloride (20 mg/kg, 500 μL; Merck, P0884) or sotalol hydrochloride (10 mg/kg, 500 μL; Merck, S0278) or clozapine N-oxide dihydrochloride (CNO water-soluble, hellobio, HB6149; 10mg/kg, 400μL) or saline (0.9% NaCl, 400/500μL) were administered by intraperitoneal injection. After the injection, the mice were isolated for thirty minutes before the beginning of the experimental procedures.

### Electrophysiology recordings

As previously described, mice were injected with inhibitory DRE-ADDs or mCherry in the BLA. The same cohort of mice was injected with undiluted AAV-5/2-hSyn1-chI-ChrimsonR_tdTomato (titer: 5.3 x 10E12 vg/ml) in the MGN. After 4-5 weeks of expression, the mice were anesthetized using 0.5mg/kg FMM with the following mixture 0.05 mg/ml of Fentanyl [(Hameln, 007007) 0.05 mg/kg], plus 5 mg/ml of Midazolam [(Hameln, 002124) 5 mg/kg] and 1 mg/ ml of Medetomidine [(VM Pharma, 087896)]. After the induction of the anesthesia, the mice were placed on the stereotaxic frame. A 32-channel optoelectrode (Poly3, Neuronexus) was placed at the following coordinates AP: 1.6 mm ML: 3.48 DV: 3.5±0.1. The neural data were amplified and digitized at 25 kHz. The input-output curve was recorded 30 minutes before and one hour after the CNO injection using a pulse of 0.5/1ms, 638nm.

At the end of the experiment, mice were sacrificed by cervical dislocation, and brains were extracted and kept in 10% formalin for 24h. Afterward, the brains were processed to confirm viral expression and electrode location.

### Fiber photometry recordings and analysis

All the recordings were performed with Doric fiber photometry system composed of a led-driver, a fiber photometry console, and a Doric mini-cube with 460-490 nm for GCaMP excitation, 415 nm for isosbestic excitation, and 580-650 nm for optical stimulation [ilFMC5-G2_IE(400-410)_E(460-490)_F(500-540)_O(580-680)_S)]. A low-autofluorescence patch cord (200 nm or 300 nm, 0.37 NA) and a pigtailed rotary joint (200 nm, 0.37 NA) were used. The latter were bleached for 5 hours before each experiment via a 473 nm laser at 15-17 mW. The GCaMP signal was amplified with a Doric amplifier with a 10x gain and recorded with Doric Neuroscience Studio software (Version 5.4.1.23) at 11 kHz. The signal was downsampled at 120 Hz for analysis. Light power at the patchcord tip was set between 30 to 35 μW for 470 nm excitation.

For synchronization with the looming stimulus presentation, a National Instrument board (NI USB 6003) was used to timestamp the looming stimulus presentation over the calcium signal. For synchronization of the tone and shock presentation, an input-output box connected to the aversive conditioning system was used.

After four to five weeks of virus expression, mice were handled for two to three days before the beginning of the experiments. Before each recording, the fiber-optic cannula was cleaned with CleanClicker (Thorlab; USA). To reduce bleaching during the behavioral experiments, the habituation to the looming stimulus and the extinction to the tone were conducted without recording the GCaMP signal.

A customized MATLAB script was used for the analysis. All the traces with a sudden change in the isosbestic signal were discarded in the final analysis.

### Immunofluorescence

The mice were anesthetized with Isofluorane and euthanized by cervical dislocation. The brains were harvested and stored for 24 hours in 10% formalin at room temperature. Then, the brains were sliced into 100-120 μm thick slices in PBS on Leica Vibratome (VT1000 S).

To visualize the extent of the lesion, the brains were stained for NeuN, and GFP Slices were permeabilized with PBS-Triton X 0.5% plus 10% of Normal Goat Serum (NGS) and blocked in 10% Bovine Goat Serum (BSA) for 90 minutes at room temperature. Subsequently, the slices were incubated with a mixture of anti-NeuN antibody mouse (Merk Millipore, MAB377; 1:500) and anti-GFP (Invitrogen, CAB4211, 1:1000) in PBS-Triton X 0.3%, 1% NGS and 5% BSA and the incubation lasted for 72 hours at 4°C. At the end of the 72h incubation, the slices were washed three times in PBS. The slices were incubated in Cyanine 3 (Cy3) goat anti-mouse (Thermo Fisher Scientific, A10521, 1:500) and Alexa Fluor 488 goat anti-rabbit (Thermo Fisher Scientific, A-11008, 1:1000) in PBS-Triton X 0.3%, 1% NGS and 5% BSA for 24 hours at 4°C. Nuclear staining was performed by using 1:1000 of DAPI (Sigma, D9542) for 30 minutes at room temperature. Brain slices were mounted on polysine glass slides with coverslips using Fluoromount G (Southern Biotech).

### Imaging and cell counting

Imaging was performed by using a virtual slide scanner (Olympus VS120, Japan). Tile images were taken by the whole brain slides by using 10X (UPLSAPO 2 10x / 0,40) or 20X objective (UPLSAPO 20x / 0,75). The emission wavelength for Alexa 488 was 518 nm with 250 ms of exposure time. For Cy3, the emission wavelength was 565 nm with 250 ms of exposure time.

GFP-positive cells were counted manually by using ImageJ. The experimenter, blind to the treatment, defined a region of interest and performed the cell counting.

### Statistics

Statistical analyses were performed by using GraphPad Prism 9. All the data are represented as mean ± SEM, and they were tested for normality using Shapiro-Wilk and D’Agostino-Pearson normality test. If the data represented a normal distribution, a parametric test was used. The statistical methods and the corresponding p-values are reported in the figure legend. All the data were screened for outliers by using ROUT test (Q=0.5%).

## Acknowledgements

We thank R. Malinow, J. Piriz, F. Ferenc Mátyás and members of the Nabavi laboratory for suggestions. We thank Jean-Charles Paterna and Viral Vector Facility (VVF) of the Neuroscience Center Zurich (ZNZ) for support. We thank Zachary Leamy for comments on the manuscript. This study was supported by Independent Research (DFF), Novo Nordisk Foundation (NNF16OC0023368), and AUFF NOVA grants to SN. Additionally, SN was supported by the Danish Research Institute of Translational Neuroscience (19958), and by PROMEMO (Center of Excellence for Proteins in Memory funded by the Danish National Research Foundation) (DNRF133). Figure were Created with BioRender.com.

## Contributions

I.F., V.K., K.Y. and S.N. designed the experiments. V.K., I.F. and N.M.J. performed the surgeries and behavioral testing. V.K. and I.F. acquired the fiber photometry data. I.F. acquired the electrophysiology data with the help of V.K. P.K. wrote a MATLAB script for processing the fiber photometry data. V.K. performed the data analyses. V.K. performed immunostainings, imaging and cell quantifications. V.K., I. F. and S.N. wrote the manuscript with input from N.M.J. V.K. made the figures. I.F. and S.N. conceptualized the project.

## Supplementary Figures

**Supplementary Figure 1.**
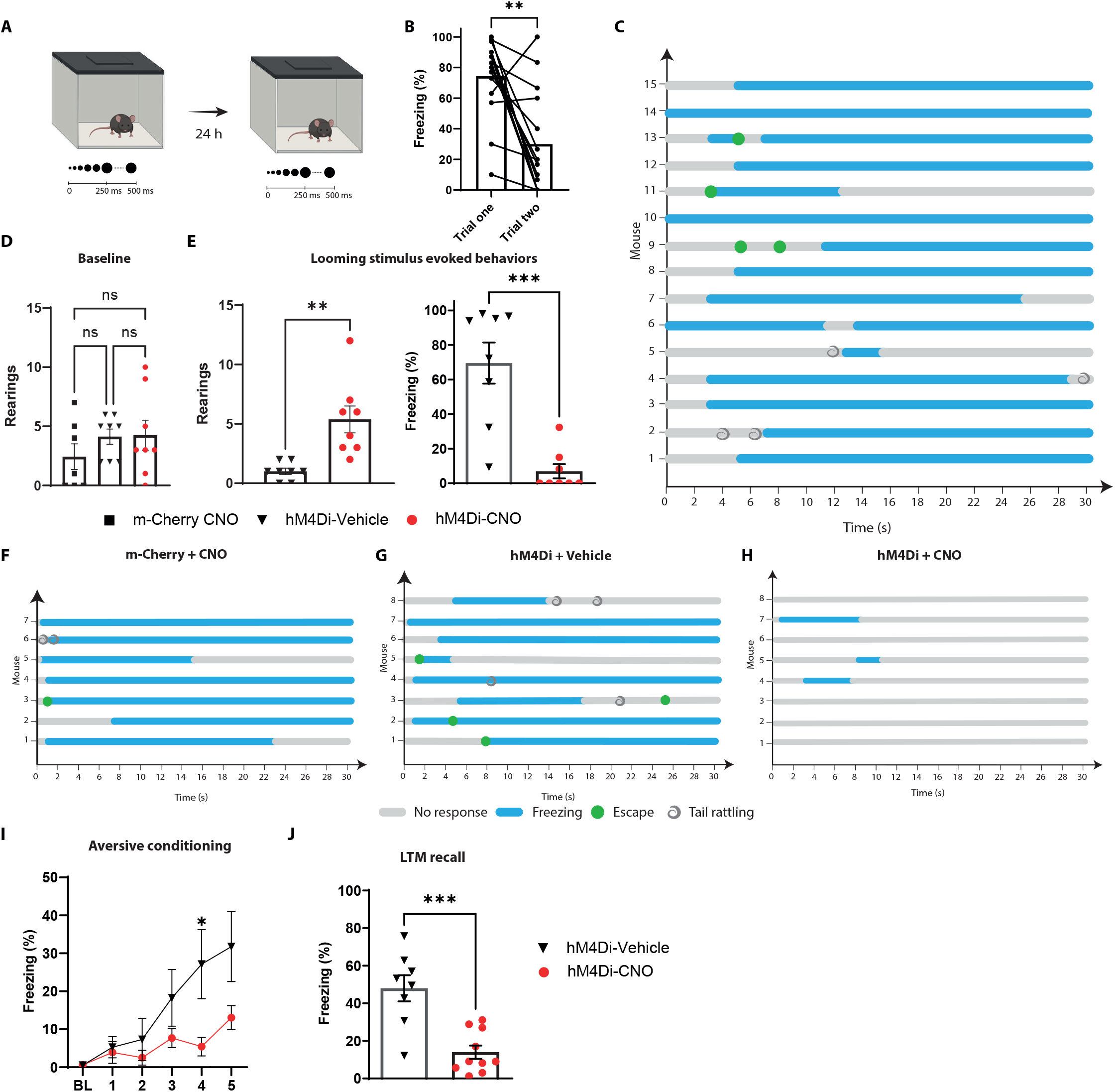
The looming stimulus response gets habituated, and it is dependent on the BLA. **A)** Timeline showing the different stages of the experiment in naïve mice. **B)** The looming-evoked freezing is reduced after several presentations of the looming stimulus (n=15, Wilcoxon test, p-value=0.0014). **C)** Ethograms showing the defensive responses for each individual during the first exposure to the looming stimulus. **D)** The number of rearing events are comparable between the three groups during the pre-looming stimulus period (Kruskal-Wallis test, F=3,23, p-value=0.3463). **E)** Right: The number of rearing events are significantly higher in the hM4Di-CNO (n=8) compared to the hM4Di-Vehicle group (n=8) during the looming stimulus exposure (Unpaired t-test, p-value=0.0021). Left: The hM4Di-CNO (n=8) group showed a significant reduction in the freezing levels when exposed to the looming stimulus compared to the hM4Di-Vehicle group (n=8, Unpaired t-test, p-value=0.0002). **F,G,H)** Ethograms show the defensive responses to the looming stimulus for the m-Cherry-CNO, hM4Di-Vehicle, and hM4Di-CNO, respectively. **I)** The graph shows the freezing levels during the baseline period (BL) and the five pairings during the conditioning protocol. The hM4Di-CNO group (n=10) showed a significant reduction in CS-evoked freezing than the hM4Di-Vehicle group (n=8; Repeated-measures ANOVA for group by time interactions, F: 5,75=3.808, p-value= 0.0040 with Sìdak test correction). **J)** The graph shows the CS-evoked freezing in a new context. The hM4Di-CNO group (n=10) showed a significant reduction in tone-evoked freezing than the hM4Di-Vehicle group (n=8; Unpaired t-test, p-value=0.0003). Data of hM4Di-CNO group are a replica as in figure 1. Results are reported as mean ± S.E.M. ns, non-significant; *, p<0.05; **, p<0.01; ***, p<0.001.

**Supplementary Figure 2.**
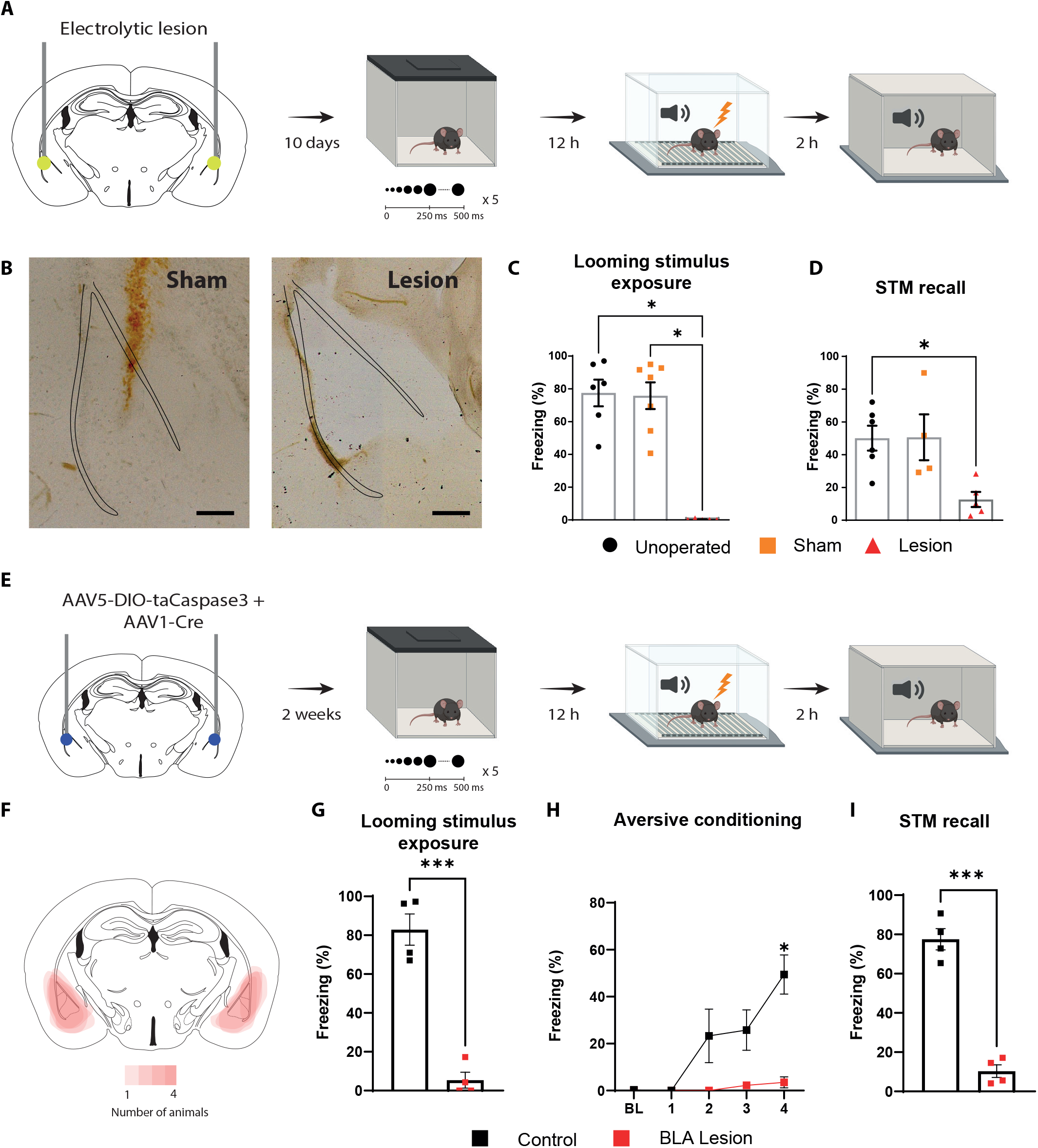
Lesions of the BLA block the defensive responses to the looming stimulus and to the aversive conditioning. **A)** Timeline showing the different stages of the experiment. **B)** Representative images showing the electrode placement in a mouse from the sham group (left) and the extent of the lesion in the BLA in a mouse from the lesioned group (right). Scale bar, 200 μm. **C)** The freezing levels are significantly reduced in the lesioned group (n=5) compared to the Unoperated (n=6) and Sham groups (n=7; Kruskal-Wallis test, F: 3,18; p-value=0.0016 with Dunn’s multiple comparison test). **D)** Tone-evoked freezing in a new context during the short-term memory recall (STM). The lesioned group (n=5) showed a significant reduction in CS-evoked freezing than the unoperated (n=6) and sham groups (n=4; Kruskal-Wallis test, F: 3,15; p-value=0.0049 with Dunn’s multiple comparison test). **E)** Timeline showing the different stages of the experiment. **F)** Overlay of the maximum extent of the lesion in the BLA-lesioned group (n=4). **G)** The looming stimulus-evoked freezing is significantly reduced in the mice from the BLA-lesioned group (n=4) compared to the control group (n=4) injected with DIO-taCaspase3 only (Unpaired t-test, p-value=0.0001). **H)** During aversive conditioning, the mice from the BLA-lesioned group (n=4) showed a significant reduction in the tone-evoked freezing compared to the control group (n=4; Repeated-measures ANOVA for group by time interactions, F: 4,24=10.02, p-value< 0.0001 with Sìdak test correction). **I)** Tone-evoked freezing in a new context during the short-term memory recall (STM). The BLA-lesioned group (n=4) showed a significant reduction in tone-evoked freezing compared to the control group (n=4; Unpaired t-test, p-value<0.0001). Results are reported as mean ± S.E.M. *, p<0.05; ***, p<0.001.

**Supplementary Figure 3.**
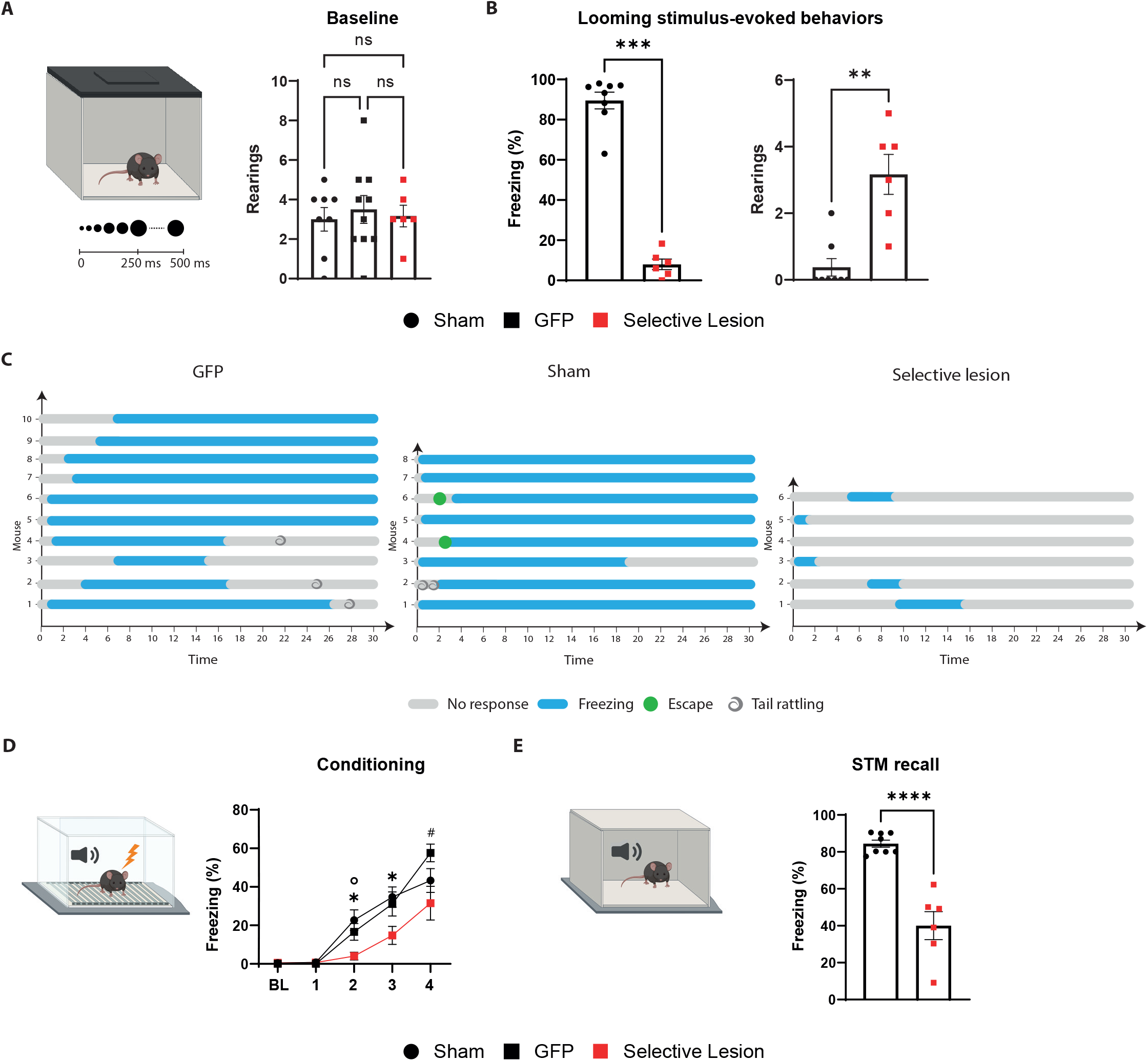
The selective lesion of the BLA-projecting neurons in MGN impairs the defensive responses to the looming stimulus and to the aversive conditioning. **A)** The number of rearing events are comparable between the three groups during the pre-looming stimulus period (Ordinary One-way ANOVA, F=1.017 (2,21), p-value=0.3787). **B)** Left: The freezing levels are significantly reduced in the selective lesion group (n=6) compared to the sham group (n=8; Mann-Whitney test, p-value=0.0007). Right: The number of rearing events is significantly higher in the selective lesion (n=6) compared to the sham group (n=8) during the looming stimulus exposure (Mann-Whitney test, p-value=0.0020). **C)** Ethograms of the defensive responses to the looming stimulus for each group. **D)** During aversive conditioning, the mice from the selective lesion group (n=6) showed a significant reduction of the tone-evoked freezing response compared to the GFP group (n=10) and sham group (n=8; Repeated-measures ANOVA for group by time interactions, F: 8,84=2.758, p-value= 0.0093 with Holm-Sìdak test correction). **E)** CS-evoked freezing in a new context during the short-term memory recall. The selective lesion group (n=6) showed a significant reduction in CS-evoked freezing compared to the sham group (n=8; Unpaired t-test, p-value<0.0001). The data for the selective lesion group are the replica of Fig. 2. The results are reported as mean ± S.E.M. ns, non-significant; #,*, p<0.05; **, p<0.01; ***, p<0.001; ****, p<0.0001.

**Supplementary Figure 4.**
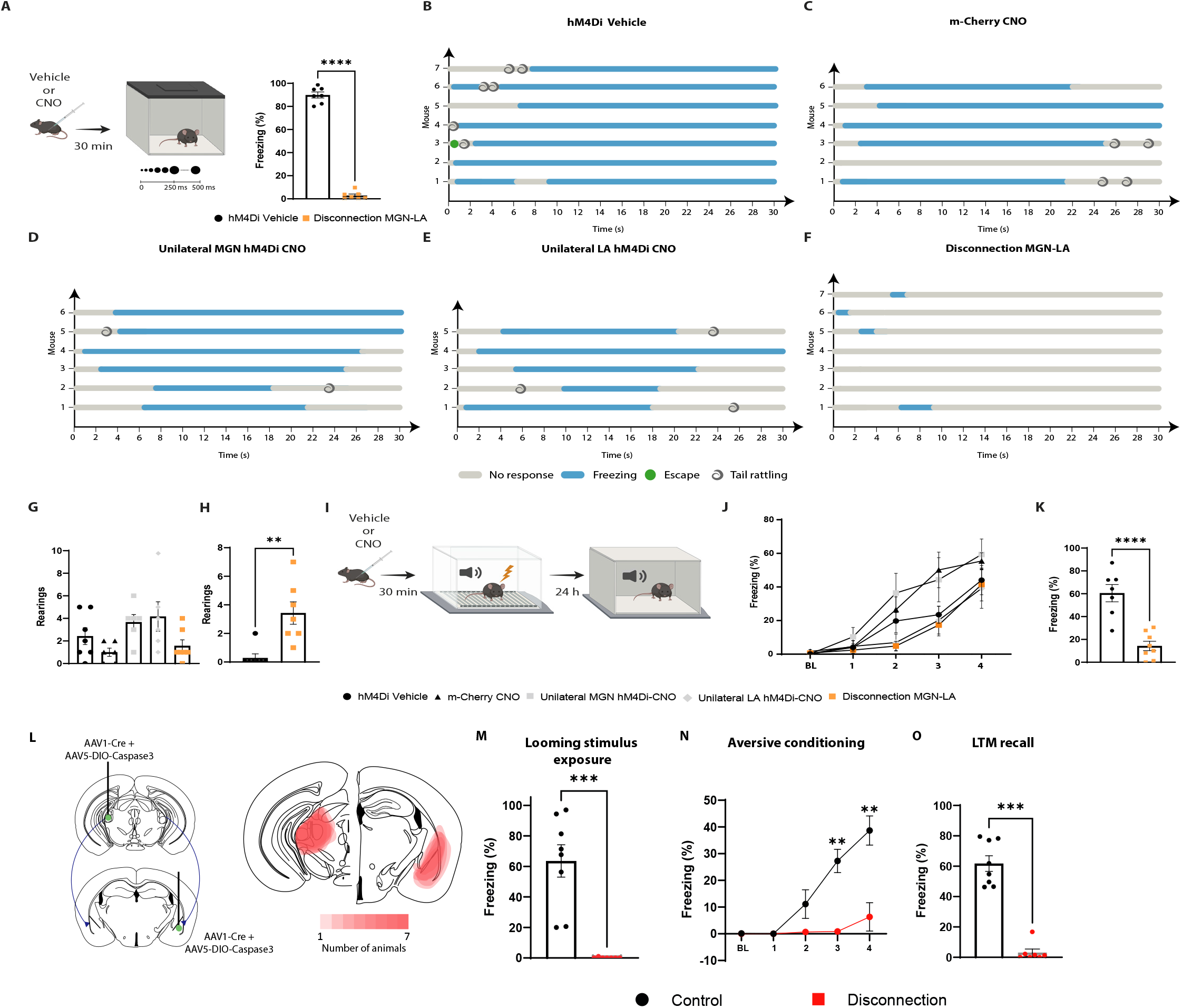
Reversible and irreversible controlateral disconnection of the MGN-BLA pathway impairs the defensive responses to the looming stimulus and to the aversive conditioning. **A)** The looming stimulus-evoked freezing is significantly reduced in the disconnection MGN-BLA group (n=7) to the hM4Di-vehicle group (n=7; Unpaired t-test, p-value<0.0001). **B,C,D,E,F)** Ethograms show the defensive responses to the looming stimulus for the hM4Di-Vehicle, m-Cherry-CNO, Unilateral MGN hM4Di-CNO, Unilateral BLA hM4Di-CNO and disconnection MGN-BLA, respectively. **G)** The number of rearing events are comparable between the five groups during the pre-looming stimulus period (Kruskal-Wallis test, F (5,32), p-value=0.0427). **H)** The number of rearing events are significantly higher in the disconnection MGN-BLA (n=7) compared to the hM4Di-Vehicle (n=7) during the looming stimulus exposure (Mann-Whitney test, p-value=0.0023). **I)** The day after the looming stimulus exposure, the mice were reinjected with CNO thirty minutes before aversive conditioning. Twenty-four hours later, the mice were exposed to the CS in a new context in a CNO-free trial. **J)** The graph shows the CS-evoked freezing during the baseline (BL) and the four pairings of the conditioning session. The contralateral disconnection of the MGN-BLA pathway did not affect the CS-evoked freezing during the conditioning session (Repeated-measures ANOVA for group by time interactions, F: 16,120=1.844, p-value=0.0328 with Tukey test correction). **K)** The disconnection MGN-BLA group (n=8) showed a significant reduction in CS-evoked freezing than the hM4Di-Vehicle group (n=7) during the recall session (Unpaired t-test, p-value<0.0001). **L)** Viral strategy and overlay of the extent of the lesion in the MGN and in the BLA. **M)** The looming stimulus-evoked freezing level is significantly reduced in the disconnection group (n=7) compared to the control group (n=8; Mann-Whitney test, p-value=0.0003) that was injected contralaterally with DIO-taCaspase3 only. **N)** The disconnection group (n=7) showed a significant reduction in CS-evoked freezing compared to the control group across the conditioning protocol (n=8; Repeated-measures ANOVA for group by time interactions, F: 4,52=10.10, p-value< 0.0001 with Sìdak test correction). **O)** The CS-evoked freezing are significantly reduced in the disconnection group (n=7) compared to the control group (n=8; Mann-Whitney test, p-value=0.0003). Data of disconnection MGN-BLA group is the same as in figure 4. Results are reported as mean ±S.E.M. *, p<0.05; **, p<0.01; ***, p<0.001; ****, p<0.0001.

**Supplementary Figure 5.**
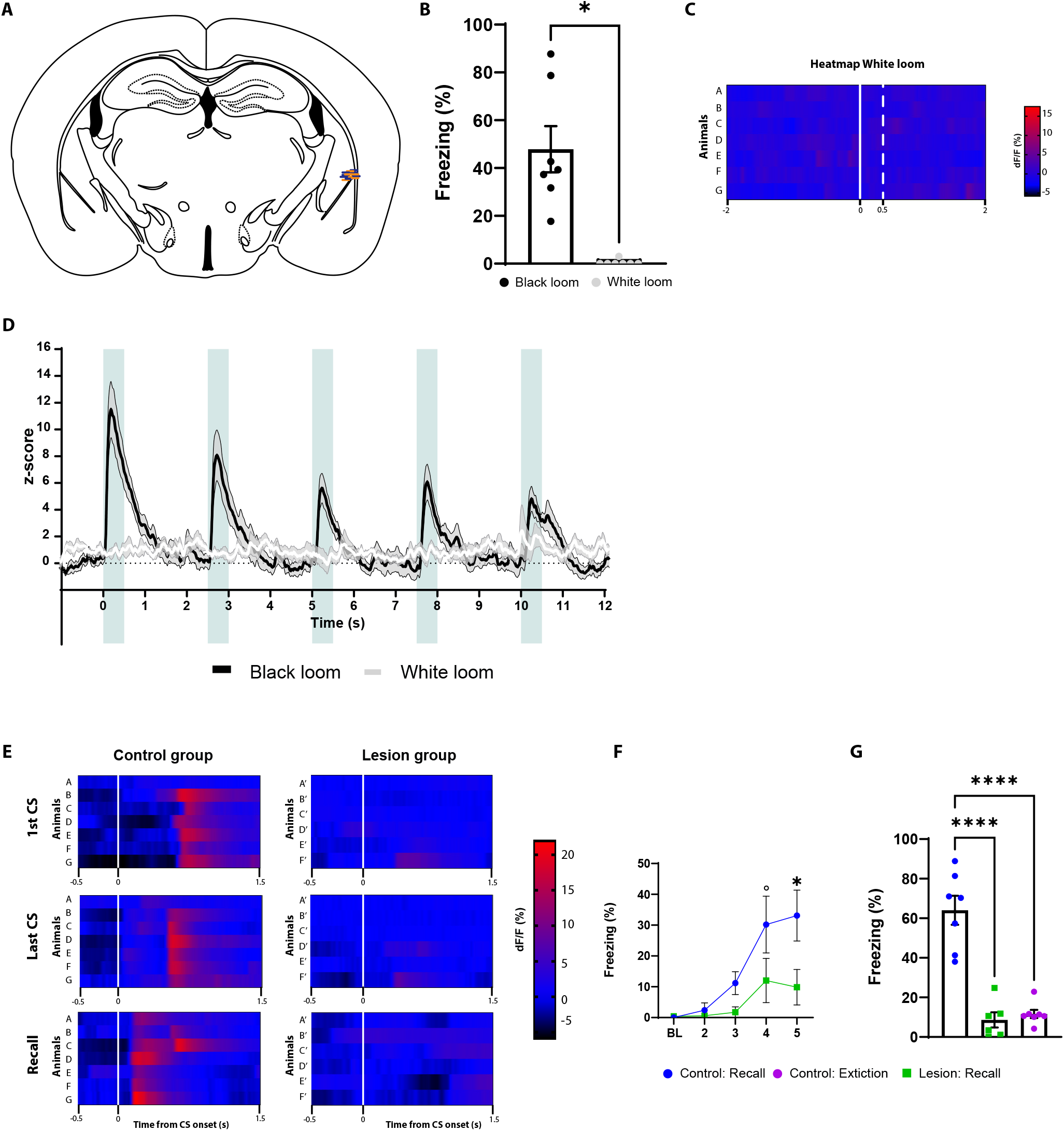
The BLA response is not activated by the white looming stimulus and the MGN lesions affect the CS processing in the BLA. **A)** Each blue and orange lines represent the optic fiber placement of an individual mouse from the control and lesion groups, respectively. **B)** The freezing level evoked by the black-looming stimulus (n=7) is significantly higher than the one evoked by the white loom (n=7, Wilcoxon test, p-value=0.0156). **C)** Heatmap representing the individual response evoked by the white looming. **D)** Z-score of the Ca2+ response during the black looming stimulus (n=7; in black), and to the white looming stimulus (n=7; in white). **E)** Heatmap representing the individual response to the first CS (top), last CS of the aversive conditioning (middle), and the recall (bottom) for the control group (left column) and the Lesion group (right column). **F)** The lesion group (n=6) showed a significant reduction in the CS-evoked freezing during the last pairing compared to the control group (n=7; Repeated-measures ANOVA for group by time interactions, F: 4,44=2.817, p-value=0.0364 with Sìdak test correction). **G)** The lesion of the MGN reduced the CS-evoked freezing during the recall session, similarly to the CS-evoked freezing level after extinction training (n=7) in the control group (Ordinary one-way ANOVA, F (2,17) = 39.15, p-value<0.0001). Results are reported as mean ± S.E.M. °, p<0.10; *, p<0.05. Data of black loom group are a replica of figure 5. Results are reported as mean ± S.E.M. *, p<0.05.

**Supplementary Figure 6.**
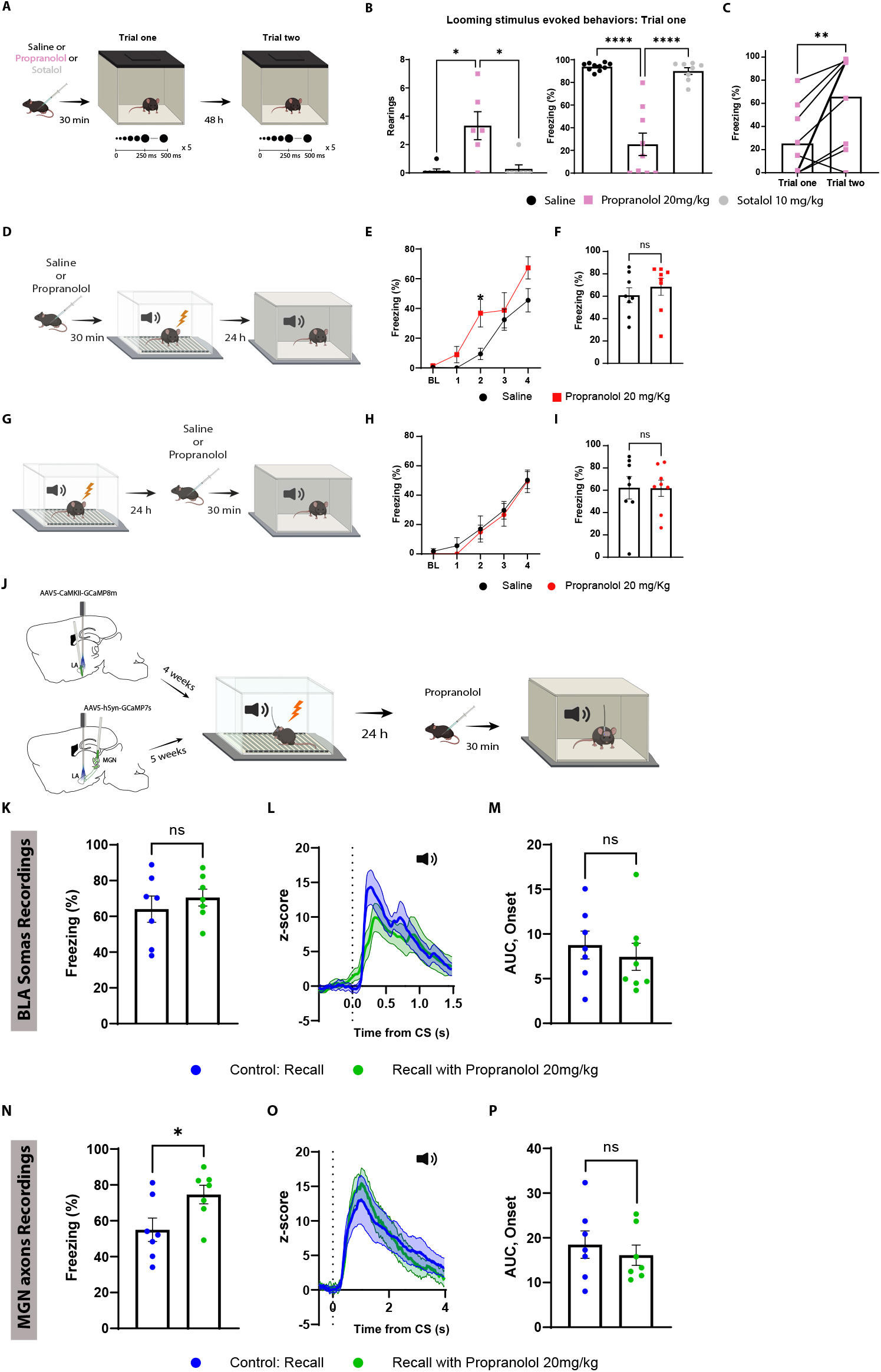
Propranolol reduces defensive response to a looming stimulus but it does not impair aversive conditioning. **A)** Diagram showing the timeline of the experiment. **B)** Left: The number of rearing events are significantly higher in the propranolol group (n=6) compared to sotalol (n=7) and saline (n=7) groups during the looming stimulus exposure (Kruskal-Wallis test, p-value=0.0026). Right: The propranolol group (n=9) showed a significant reduction in the freezing levels compared to the saline group (n=10) and the sotalol group (n=8; Ordinary one-way ANOVA, F (2,24) = 41.89, p-value<0.0001). **C)** The propranolol group showed a significant increase in the freezing levels evoked by the looming stimulus during the re-exposure trial (Paired t-test, p-value=0.0053). **D)** Timeline showing the different stages of the experiment. **E)** The freezing levels during the baseline (BL) period and the consecutive four pairs are similar between the saline (n=8) and the propranolol (n=8) groups. The only significant difference is in the CS-evoked freezing levels during the second pairing of the conditioning (Two-way ANOVA, F (4, 56)= 2.084, p-value= 0.0950). **F)** There is no difference in the CS-evoked freezing levels between the saline (n=8) and the propranolol group (n=8; Mann-Whitney test, p-values=0.4866). **G)** A separate cohort of mice was conditioned and injected with saline or propranolol thirty minutes before long term memory recall in a new context. **H)** The freezing levels during the BL period and the consecutive four pairs are comparable between the saline (n=8) and the propranolol (n=8) groups (Two-way ANOVA, F (4, 70)= 0.04946, p-value= 0.9953). **I)** There is no difference in the CS-evoked freezing levels between the saline (n=8) and the propranolol group (n=8) during the recall session (Unpaired t-test, p-values=0.9732). **J)** Timeline showing the different stages of the experiments. **K)** The CS-evoked freezing is comparable between a group of mice injected with propranolol (n=7) compared to the control group (n=7; Unpaired t-test, p-value= 0.4732). The control group is a replica of Figure S6. **L)** Z-score of the Ca2+ responses in the BLA to the first CS presentation during the recall for the control group (n=7) and the propranolol injected group (n=7) in the group injected with propranolol before the recall session. The control group is a replica of Figure 5. **M)** The AUC is unchanged between the controls (n=7) and the group injected with propranolol (n=7; Unpaired t-test, p-value=0.5535). **N)** CS-evoked freezing is significantly increased in the propranolol injected group (n=7) compared to the control group (n=7; Unpaired t-test, p-value= 0.0359). **O)** Z-score of the Ca2+ responses in the MGN axons to the first CS presentation during the recall for the control group (n=7) and the propranolol injected group (n=7) in the group injected with propranolol before the recall session. The control group is a replica of Figure 3. **P)** The AUC is unchanged between the controls (n=7) and the group injected with propranolol (n=7; Unpaired t-test, p-value=0.5456). Results are reported as mean ± S.E.M. ns, non-significant; *, p<0.05; **, p<0.01; ****, p<0.0001.

**Supplementary Figure 7.**
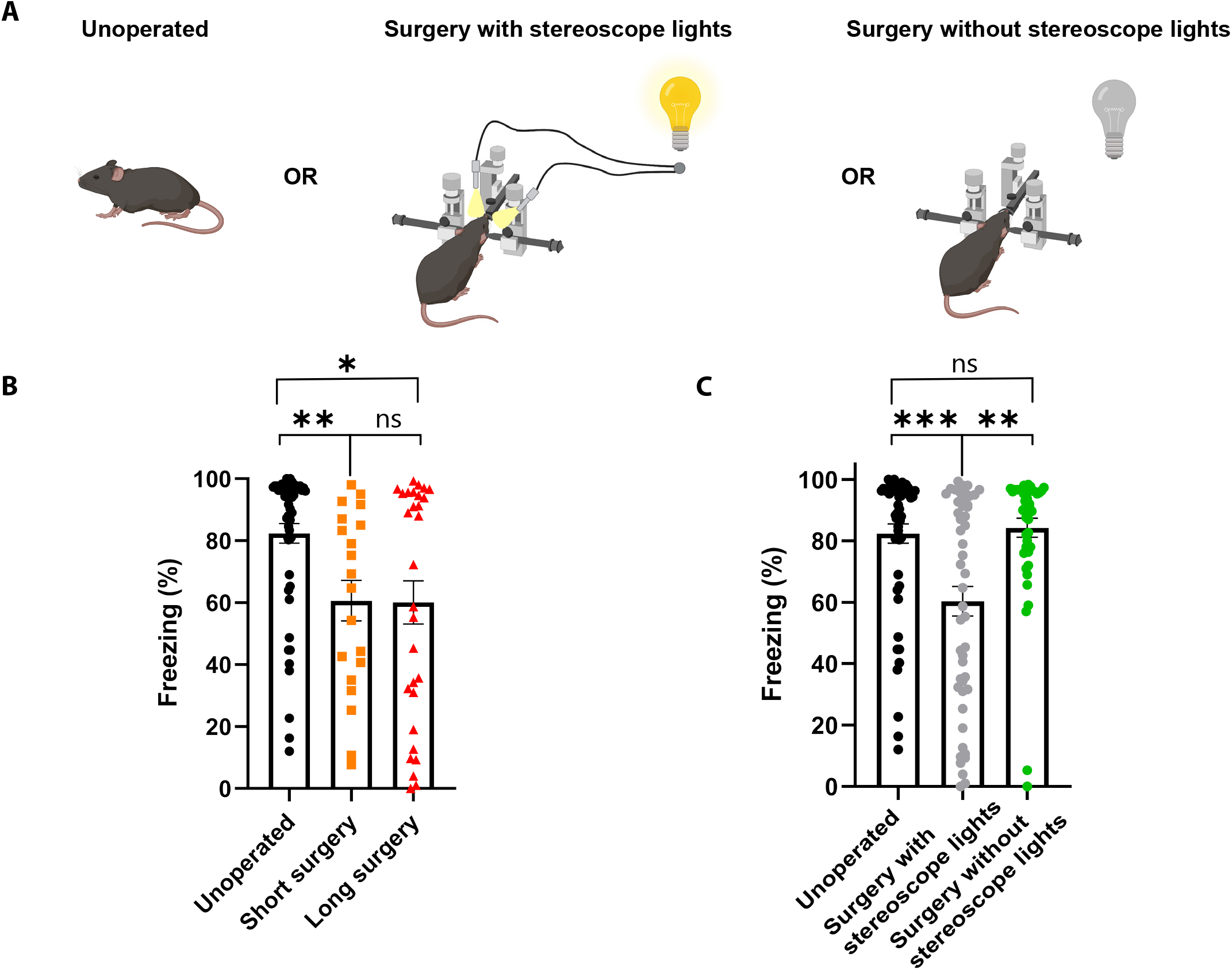
Effect of the surgery on the looming stimulus-evoked responses. **A)** Diagram of the three experimental groups. **B)** The surgery procedures significantly reduced the looming stimulus-evoked freezing regardless of the length of the surgeries (Unoperated, n=54; Short surgery, n=20; Long surgery, n=29; KruskalWallis test, p-value=0.0007). **C)** Surgery without the stereoscope lights group (n=45) showed no significant difference in the looming stimulus-evoked freezing compared to the unoperated group (n=54). Short- and Long-surgery groups were pooled together in the group Surgery with stereoscope lights (n=49; Kruskal-Wallis test, p-value=0.0002). Results are reported as mean ±S.E.M. ns, not significant; *, p<0.05; **, p<0.01; ***, p<0.001.

## References

Anglada-Figueroa, D., & Quirk, G. J. (2005). Lesions of the basal amygdala block expression of conditioned fear but not extinction. Journal of Neuroscience, 25(42), 9680–9685. https://doi.org/10.1523/JNEUROSCI.2600-05.2005

Armbruster, B. N., Li, X., Pausch, M. H., Herlitze, S., & Roth, B. L. (2007). Evolving the lock to fit the key to create a family of G protein-coupled receptors potently activated by an inert ligand. Proceedings of the National Academy of Sciences of the United States of America, 104(12), 5163–5168. https://doi.org/10.1073/pnas.0700293104

Barbano, M. F., Wang, H.-L., Zhang, S., Miranda-Barrientos, J., Estrin, D. J., Figueroa-González, A., Liu, B., Barker, D. J., & Morales, M. (2020). VTA Glutamatergic Neurons Mediate Innate Defensive Behaviors. Neuron, 0(0), 1–15. https://doi.org/10.1016/j.neuron.2020.04.024

Barker, G. R. I., Banks, P. J., Scott, H., Ralph, G. S., Mitrophanous, K. A., Wong, L. F., Bashir, Z. I., Uney, J. B., & Warburton, E. C. (2017). Separate elements of episodic memory subserved by distinct hippocampal-prefrontal connections. Nature Neuroscience, 20(2), 242–250. https://doi.org/10.1038/nn.4472

Barsy, B., Kocsis, K., Magyar, A., Babiczky, Á., Szabó, M., Veres, J. M., Hillier, D., Ulbert, I., Yizhar, O., & Mátyás, F. (2020). Associative and plastic thalamic signaling to the lateral amygdala controls fear behavior. Nature Neuroscience, 23(5), 625–637. https://doi.org/10.1038/s41593-020-0620-z

Bindi, R. P., Baldo, M. V. C., & Canteras, N. S. (2018). Roles of the anterior basolateral amygdalar nucleus during exposure to a live predator and to a predator-associated context. Behavioural Brain Research, 342(October 2017), 51–56. https://doi.org/10.1016/j.bbr.2018.01.016

Blair, H. T., Schafe, G. E., Bauer, E. P., Rodrigues, S. M., & LeDoux, J. E. (2001). Synaptic plasticity in the lateral amygdala: A cellular hypothesis of fear conditioning. Learning and Memory, 8(5), 229–242. https://doi.org/10.1101/lm.30901

Blanchard, D. C., Griebel, G., & Blanchard, R. J. (2001). Mouse defensive behaviors: pharmacological and behavioral assays for anxiety and panic. Neuroscience & Biobehavioral Reviews, 25(3), 205–218. https://doi.org/10.1016/S0149-7634(01)00009-4

Bordi, F., & LeDoux, J. (1992). Sensory tuning beyond the sensory system: an initial analysis of auditory response properties of neurons in the lateral amygdaloid nucleus and overlying areas of the striatum. The Journal of Neuroscience, 12(7), 2493. https://doi.org/10.1523/JNEUROSCI.12-07-02493.1992

Bordi, Fabio, LeDoux, J., Clugnet, M. C., & Pavlides, C. (1993). Single-Unit Activity in the Lateral Nucleus of the Amygdala and Overlying Areas of the Striatum in Freely Behaving Rats: Rates, Discharge Patterns, and Responses to Acoustic Stimuli. Behavioral Neuroscience, 107(5), 757–769. https://doi.org/10.1037/0735-7044.107.5.757

Bordi, Fabio, & LeDoux, J. E. (1994). Response properties of single units in areas of rat auditory thalamus that project to the amygdala - II. Cells receiving convergent auditory and somatosensory inputs and cells antidromically activated by amygdala stimulation. Experimental Brain Research, 98(2), 275–286. https://doi.org/10.1007/BF00228415

Campeau, S., & Davis, M. (1995). Involvement of subcortical and cortical afferents to the lateral nucleus of the amygdala in fear conditioning measured with fear-potentiated startle in rats trained concurrently with auditory and visual conditioned stimuli. Journal of Neuroscience, 15(3), 2312–2327. https://doi.org/10.1523/JNEU-ROSCI.15-03-02312.1995

Carr, J. A. (2015). I’ll take the low road: The evolutionary underpinnings of visually triggered fear. Frontiers in Neuroscience, 9(OCT), 1–13. https://doi.org/10.3389/fnins.2015.00414

Dana, H., Sun, Y., Mohar, B., Hulse, B. K., Kerlin, A. M., Hasseman, J. P., Tsegaye, G., Tsang, A., Wong, A., Patel, R., Macklin, J. J., Chen, Y., Konnerth, A., Jayaraman, V., Looger, L. L., Schreiter, E. R., Svoboda, K., & Kim, D. S. (2019). High-performance calcium sensors for imaging activity in neuronal populations and microcompartments. Nature Methods 2019 16:7, 16(7), 649–657. https://doi.org/10.1038/s41592-019-0435-6

Do Monte, F. H. M., Canteras, N. S., Fernandes, D., Assreuy, J., & Carobrez, A. P. (2008). New perspectives on β-adrenergic mediation of innate and learned fear responses to predator odor. Journal of Neuroscience, 28(49), 13296–13302. https://doi.org/10.1523/JNEUROSCI.2843-08.2008

Edeline, J. M., & Weinberger, N. M. (1992). Associative Retuning in the Thalamic Source of Input to the Amygdala and Auditory Cortex: Receptive Field Plasticity in the Medial Division of the Medial Geniculate Body. Behavioral Neuroscience, 106(1), 81–105. https://doi.org/10.1037/0735-7044.106.1.81

Eldridge, M. A. G., Lerchner, W., Saunders, R. C., Kaneko, H., Krausz, K. W., Gonzalez, F. J., Ji, B., Higuchi, M., Minamimoto, T., & Richmond, B. J. (2015). Chemogenetic disconnection of monkey orbitofrontal and rhinal cortex reversibly disrupts reward value. Nature Neuroscience 2016 19:1, 19(1), 37–39. https://doi.org/10.1038/nn.4192

Evans, D. A., Stempel, A. V., Vale, R., & Branco, T. (2019). Cognitive Control of Escape Behaviour. Trends in Cognitive Sciences, 23(4), 334–348. https://doi.org/10.1016/j.tics.2019.01.012

Evans, D. A., Stempel, A. V., Vale, R., Ruehle, S., Lefler, Y., & Branco, T. (2018). A synaptic threshold mechanism for computing escape decisions. Nature, 558(7711), 590–594. https://doi.org/10.1038/s41586-018-0244-6

Fadok, J. P., Krabbe, S., Markovic, M., Courtin, J., Xu, C., Massi, L., Botta, P., Bylund, K., Müller, C., Kovacevic, A., Tovote, P., & Lüthi, A. (2017). A competitive inhibitory circuit for selection of active and passive fear responses. Nature, 542(7639), 96–99. https://doi.org/10.1038/nature21047

Grewe, B. F., Gründemann, J., Kitch, L. J., Lecoq, J. A., Parker, J. G., Marshall, J. D., Larkin, M. C., Jercog, P. E., Grenier, F., Li, J. Z., Lüthi, A., & Schnitzer, M. J. (2017). Neural ensemble dynamics underlying a long-term associative memory. Nature, 543(7647), 670–675. https://doi.org/10.1038/nature21682

Gross, C. T., & Canteras, N. S. (2012). The many paths to fear. Nature Reviews Neuroscience, 13(9), 651–658. https://doi.org/10.1038/nrn3301

Hayley, S., Borowski, T., Merali, Z., & Anisman, H. (2001). Central monoamine activity in genetically distinct strains of mice following a psychogenic stressor: Effects of predator exposure. Brain Research, 892(2), 293–300. https://doi.org/10.1016/S0006-8993(00)03262-5

Headley, D. B., Kanta, V., Kyriazi, P., & Paré, D. (2019). Embracing Complexity in Defensive Networks. Neuron, 103(2), 189–201. https://doi.org/10.1016/j.neuron.2019.05.024

Hu, H., Real, E., Takamiya, K., Kang, M. G., Ledoux, J., Huganir, R. L., & Malinow, R. (2007). Emotion Enhances Learning via Norepinephrine Regulation of AMPA-Receptor Trafficking. Cell, 131(1), 160–173. https://doi.org/10.1016/j.cell.2007.09.017

Isosaka, T., Matsuo, T., Yamaguchi, T., Funabiki, K., Nakanishi, S., Kobayakawa, R., & Kobayakawa, K. (2015). Htr2a-Expressing Cells in the Central Amygdala Control the Hierarchy between Innate and Learned Fear. Cell, 163(5), 1153–1164. https://doi.org/10.1016/j.cell.2015.10.047

Iwata, J., LeDoux, J. E., Meeley, M. P., Arneric, S., & Reis, D. J. (1986). Intrinsic neurons in the amygdaloid field projected to by the medial geniculate body mediate emotional responses conditioned to acoustic stimuli. Brain Research, 383(1–2), 195–214. https://doi.org/10.1016/0006-8993(86)90020-X

Janak, P. H., & Tye, K. M. (2015). From circuits to behaviour in the amygdala. In Nature (Vol. 517, Issue 7534, pp. 284–292). Nature Publishing Group. https://doi.org/10.1038/nature14188

Johansen, J. P., Cain, C. K., Ostroff, L. E., & Ledoux, J. E. (2011). Molecular mechanisms of fear learning and memory. In Cell (Vol. 147, Issue 3, pp. 509–524). NIH Public Access. https://doi.org/10.1016/j.cell.2011.10.009

Johansen, J. P., Diaz-Mataix, L., Hamanaka, H., Ozawa, T., Ycu, E., Koivumaa, J., Kumar, A., Hou, M., Deisseroth, K., Boyden, E. S., & LeDoux, J. E. (2014). Hebbian and neuromodulatory mechanisms interact to trigger associative memory formation. Proceedings of the National Academy of Sciences of the United States of America, 111(51), E5584–E5592. https://doi.org/10.1073/pnas.1421304111

Kang, S. J., Liu, S., Ye, M., Kim, D. Il, Pao, G. M., Copits, B. A., Roberts, B. Z., Lee, K. F., Bruchas, M. R., & Han, S. (2022). A central alarm system that gates multi-sensory innate threat cues to the amygdala. Cell Reports, 40(7), 111222. https://doi.org/10.1016/J.CELREP.2022.111222

Klyachko, V. A., & Stevens, C. F. (2003). Connectivity optimization and the positioning of cortical areas. Proceedings of the National Academy of Sciences of the United States of America, 100(13), 7937–7941. https://doi.org/10.1073/PNAS.0932745100/AS-SET/9513F6D4-BE3C-47E7-87DB-94B2186866D2/ASSETS/GRAPHIC/PQ1332745004.JPEG

Lecca, S., Meye, F. J., Trusel, M., Tchenio, A., Harris, J., Schwarz, M. K., Burdakov, D., Georges, F., & Mameli, M. (2017). Aversive stimuli drive hypothalamus-to-habenula excitation to promote escape behavior. ELife, 6, 1–16. https://doi.org/10.7554/eLife.30697

Lecca, S., Namboodiri, V. M. K., Restivo, L., Gervasi, N., Pillolla, G., Stuber, G. D., & Mameli, M. (2020). Heterogeneous Habenular Neuronal Ensembles during Selection of Defensive Behaviors. Cell Reports, 31(10), 107752. https://doi.org/10.1016/J.CEL-REP.2020.107752

LeDoux, J. E., Sakaguchi, A., & Reis, D. J. (1984). Subcortical efferent projections of the medial geniculate nucleus mediate emotional responses conditioned to acoustic stimuli. Journal of Neuroscience, 4(3), 683–698. https://doi.org/10.1523/JNEUROSCI.04-03-00683.1984

LeDoux, Joseph E., Farb, C., & Ruggiero, D. A. (1990). Topographic organization of neurons in the acoustic thalamus that project to the amygdala. Journal of Neuroscience, 10(4), 1043–1054. https://doi.org/10.1523/jneurosci.10-04-01043.1990

LeDoux, Joseph E., Iwata, J., Pearl, D., & Reis, D. J. (1986). Disruption of auditory but not visual learning by destruction of intrinsic neurons in the rat medial geniculate body. Brain Research, 371(2), 395–399. https://doi.org/10.1016/0006-8993(86)90383-5

Lee, Y., Oh, J. P., & Han, J. H. (2021). Dissociated role of thalamic and cortical input to the lateral amygdala for consolidation of longterm fear memory. Journal of Neuroscience, 34(3), 9561–9570. https://doi.org/10.1523/JNEUROSCI.1167-21.2021

Linke, R., De Lima, A. D., Schwegler, H., & Pape, H. C. (1999). Direct synaptic connections of axons from superior colliculus with identified thalamo-amygdaloid projection neurons in the rat: Possible substrates of a subcortical visual pathway to the amygdala. Journal of Comparative Neurology, 403(2), 158–170. https://doi.org/10.1002/(SICI)1096-9861(19990111)403:2<158::AID-CNE2>3.0.CO;2-6

Linke, Rüdiger. (1999). Differential projection patterns of superior and inferior collicular neurons onto posterior paralaminar nuclei of the thalamus surrounding the medial geniculate body in the rat. European Journal of Neuroscience, 11(1), 187–203. https://doi.org/10.1046/j.1460-9568.1999.00422.x

Liu, Y., Formisano, L., Savtchouk, I., Takayasu, Y., Szabó, G., Zukin, R. S., & Liu, S. J. (2010). A single fear-inducing stimulus induces a transcription-dependent switch in synaptic AMPAR phenotype. Nature Neuroscience, 13(2), 223–231. https://doi.org/10.1038/nn.2474

Maren, S. (1999). Neurotoxic basolateral amygdala lesions impair learning and memory but not the performance of conditional fear in rats. Journal of Neuroscience, 19(19), 8696–8703. https://doi.org/10.1523/JNEUROSCI.19-19-08696.1999

Maren, S., Aharonov, G., & Fanselow, M. S. (1996). Retrograde abolition of conditional fear after excitotoxic lesions in the basolateral amygdala of rats: Absence of a temporal gradient. Behavioral Neuroscience, 110(4), 718. https://doi.org/10.1037/0735-7044.110.4.718

Martinez, R. C., Carvalho-Netto, E. F., Ribeiro-Barbosa, É. R., Baldo, M. V. C., & Canteras, N. S. (2011). Amygdalar roles during exposure to a live predator and to a predator-associated context. Neuroscience, 172, 314–328. https://doi.org/10.1016/j.neuroscience.2010.10.033

Matsumoto, M., & Hikosaka, O. (2007). Lateral habenula as a source of negative reward signals in dopamine neurons. Nature 2007 447:7148, 447(7148), 1111–1115. https://doi.org/10.1038/nature05860

McFadyen, J., Dolan, R. J., & Garrido, M. I. (2020). The influence of subcortical shortcuts on disordered sensory and cognitive processing. Nature Reviews Neuroscience, 21(5), 264–276. https://doi.org/10.1038/s41583-020-0287-1

Mondoloni, S., Mameli, M., & Congiu, M. (2022). Reward and aversion encoding in the lateral habenula for innate and learned behaviours. Translational Psychiatry, 12(1). https://doi.org/10.1038/S41398-021-01774-0

Mongeau, R., Miller, G. A., Chiang, E., & Anderson, D. J. (2003). Neural correlates of competing fear behaviors evoked by an innately aversive stimulus. Journal of Neuroscience, 23(9), 3855–3868. https://doi.org/10.1523/jneurosci.23-09-03855.2003

Orsini, C. A., & Maren, S. (2012). Neural and cellular mechanisms of fear and extinction memory formation. Neuroscience & Biobehavioral Reviews, 36(7), 1773–1802. https://doi.org/10.1016/J.NEUBIOREV.2011.12.014

Pape, H. C., & Pare, D. (2010). Plastic synaptic networks of the amygdala for the acquisition, expression, and extinction of conditioned fear. In Physiological Reviews (Vol. 90, Issue 2, pp. 419–463). American Physiological Society. https://doi.org/10.1152/phys-rev.00037.2009

Pereira, A. G., & Moita, M. A. (2016). Is there anybody out there? Neural circuits of threat detection in vertebrates. Current Opinion in Neurobiology, 41, 179–187. https://doi.org/10.1016/j.conb.2016.09.011

Pessoa, L. (2008). On the relationship between emotion and cognition. Nature Reviews Neuroscience 2008 9:2, 9(2), 148–158. https://doi.org/10.1038/nrn2317

Pessoa, L., & Adolphs, R. (2010). Emotion processing and the amygdala: from a “low road” to “many roads” of evaluating biological significance. Nature Reviews Neuroscience 2010 11:11, 11(11), 773–782. https://doi.org/10.1038/nrn2920

Quirk, G. J., Repa, J. C., & LeDoux, J. E. (1995). Fear conditioning enhances short-latency auditory responses of lateral amygdala neurons: Parallel recordings in the freely behaving rat. Neuron, 15(5), 1029–1039. https://doi.org/10.1016/0896-6273(95)90092-6

Rodriguez-Romaguera, J., Sotres-Bayon, F., Mueller, D., & Quirk, G. J. (2009). Systemic Propranolol Acts Centrally to Reduce Conditioned Fear in Rats Without Impairing Extinction. Biological Psychiatry, 65(10), 887–892. https://doi.org/10.1016/J.BIO-PSYCH.2009.01.009

Rogan, M. T., Staubli, U. V., & LeDoux, J. E. (1997). Fear conditioning induces associative long-term potentiation in the amygdala. Nature 1997 390:6660, 390(6660), 604–607. https://doi.org/10.1038/37601

Rogan, Michael T., & LeDoux, J. E. (1995). LTP is accompanied by commensurate enhancement of auditory-evoked responses in a fear conditioning circuit. Neuron, 15(1), 127–136. https://doi.org/10.1016/0896-6273(95)90070-5

Rogan, Michael T., Leon, K. S., Perez, D. L., & Kandel, E. R. (2005). Distinct neural signatures for safety and danger in the amygdala and striatum of the mouse. Neuron, 46(2), 309–320. https://doi.org/10.1016/j.neuron.2005.02.017

Romanski, L. M., & LeDoux, J. E. (1992). Equipotentiality of thala-mo-amygdala and thalamo-cortico-amygdala circuits in auditory fear conditioning. Journal of Neuroscience, 12(11), 4501–4509. https://doi.org/10.1523/jneurosci.12-11-04501.1992

Romanski, Lizabeth M., Clugnet, M. C., Bordi, F., & LeDoux, J. E. (1993). Somatosensory and Auditory Convergence in the Lateral Nucleus of the Amygdala. Behavioral Neuroscience, 107(3), 444–450. https://doi.org/10.1037/0735-7044.107.3.444

Romanski, Lizabeth M., & LeDoux, J. E. (1992). Bilateral destruction of neocortical and perirhinal projection targets of the acoustic thalamus does not disrupt auditory fear conditioning. Neuroscience Letters, 142(2), 228–232. https://doi.org/10.1016/0304-3940(92)90379-L

Root, D. H., Mejias-Aponte, C. A., Qi, J., & Morales, M. (2014). Role of Glutamatergic Projections from Ventral Tegmental Area to Lateral Habenula in Aversive Conditioning. Journal of Neuroscience, 34(42), 13906–13910. https://doi.org/10.1523/JNEUROS-CI.2029-14.2014

Sachella, T. E., Ihidoype, M. R., Proulx, C. D., Pafundo, D. E., Medina, J. H., Mendez, P., & Piriz, J. (2022). A novel role for the lateral habenula in fear learning. Neuropsychopharmacology 2022 47:6, 47(6), 1210–1219. https://doi.org/10.1038/s41386-022-01294-5

Salay, L. D., Ishiko, N., & Huberman, A. D. (2018). A midline thalamic circuit determines reactions to visual threat. Nature, 557(7704). https://doi.org/10.1038/s41586-018-0078-2

Shang, C., Chen, Z., Liu, A., Li, Y., Zhang, J., Qu, B., Yan, F., Zhang, Y., Liu, W., Liu, Z., Guo, X., Li, D., Wang, Y., & Cao, P. (2018). Divergent midbrain circuits orchestrate escape and freezing responses to looming stimuli in mice. Nature Communications, 9(1). https://doi.org/10.1038/s41467-018-03580-7

Shang, C., Liu, Z., Chen, Z., Shi, Y., Wang, Q., Liu, S., Li, D., & Cao, P. (2015). A parvalbumin-positive excitatory visual pathway to trigger fear responses in mice. Science, 348(6242), 1472–1477. https://doi.org/10.1126/science.aaa8694

Shukla, A., & Chattarji, S. (2021). Stressed rats fail to exhibit avoidance reactions to innately aversive social calls. Neuropsychopharmacology 2021 47:6, 47(6), 1145–1155. https://doi.org/10.1038/s41386-021-01230-z

Silva, B. A., Gross, C. T., & Gräff, J. (2016). The neural circuits of innate fear: Detection, integration, action, and memorization. Learning and Memory, 23(10), 544–555. https://doi.org/10.1101/lm.042812.116

Stachniak, T. J., Ghosh, A., & Sternson, S. M. (2014). Chemogenetic Synaptic Silencing of Neural Circuits Localizes a Hypothalamus→Midbrain Pathway for Feeding Behavior. Neuron, 82(4), 797808. https://doi.org/10.1016/J.NEURON.2014.04.008

Stevens, C. F. (2012). Brain Organization: Wiring Economy Works for the Large and Small. Current Biology, 22(1), R24–R25. https://doi.org/10.1016/J.CUB.2011.11.036

Taylor, J. A., Hasegawa, M., Benoit, C. M., Freire, J. A., Theodore, M., Ganea, D. A., Innocenti, S. M., Lu, T., & Gründemann, J. (2021). Single cell plasticity and population coding stability in auditory thalamus upon associative learning. Nature Communications, 12(1), 1–14. https://doi.org/10.1038/s41467-021-22421-8

Tierney, A. J. (1986). The evolution of learned and innate behavior: Contributions from genetics and neurobiology to a theory of behavioral evolution. Animal Learning & Behavior 1986 14:4, 14(4), 339–348. https://doi.org/10.3758/BF03200077

Torromino, G., Autore, L., Khalil, V., Mastrorilli, V., Griguoli, M., Pignataro, A., Centofante, E., Biasini, G. M., De Turris, V., Ammas-sari-Teule, M., Rinaldi, A., & Mele, A. (2019). Offline ventral subiculum-ventral striatum serial communication is required for spatial memory consolidation. Nature Communications 2019 10:1, 10(1), 1–9. https://doi.org/10.1038/s41467-019-13703-3

Tovote, P., Esposito, M. S., Botta, P., Chaudun, F., Fadok, J. P., Markovic, M., Wolff, S. B. E., Ramakrishnan, C., Fenno, L., Deisseroth, K., Herry, C., Arber, S., & Lüthi, A. (2016). Midbrain circuits for defensive behaviour. Nature, 534(7606), 206–212. https://doi.org/10.1038/nature17996

Tye, K. M., Stuber, G. D., De Ridder, B., Bonci, A., & Janak, P. H. (2008). Rapid strengthening of thalamo-amygdala synapses mediates cue–reward learning. Nature 2008 453:7199, 453(7199), 1253–1257. https://doi.org/10.1038/nature06963

Wei, P., Liu, N., Zhang, Z., Liu, X., Tang, Y., He, X., Wu, B., Zhou, Z., Liu, Y., Li, J., Zhang, Y., Zhou, X., Xu, L., Chen, L., Bi, G., Hu, X., Xu, F., & Wang, L. (2015). Processing of visually evoked innate fear by a non-canonical thalamic pathway. Nature Communications, 6, 1–12. https://doi.org/10.1038/ncomms7756

Weinberger, N. M. (2011). The medial geniculate, not the amygdala, as the root of auditory fear conditioning. Hearing Research, 274(1–2), 61–74. https://doi.org/10.1016/j.heares.2010.03.093

Yang, C. F., Chiang, M. C., Gray, D. C., Prabhakaran, M., Alvarado, M., Juntti, S. A., Unger, E. K., Wells, J. A., & Shah, N. M. (2013). Sexually dimorphic neurons in the ventromedial hypothalamus govern mating in both sexes and aggression in males. Cell, 153(4), 896–909. https://doi.org/10.1016/j.cell.2013.04.017

Yilmaz, M., & Meister, M. (2013). Rapid innate defensive responses of mice to looming visual stimuli. Current Biology, 23(20), 2011–2015. https://doi.org/10.1016/j.cub.2013.08.015

Zhang, Y., Rózsa, M., Liang, Y., Bushey, D., Wei, Z., Zheng, J., Reep, D., Broussard, G. J., Tsang, A., Tsegaye, G., Narayan, S., Obara, C. J., Lim, J.-X., Patel, R., Zhang, R., Ahrens, M. B., Turner, G. C., Wang, S. S.-H., Korff, W. L.,… Looger, L. L. (2021). Fast and sensitive GCaMP calcium indicators for imaging neural populations. BioRxiv, 2021.11.08.467793. https://doi.org/10.1101/2021.11.08.467793

Zhou, Z., Liu, X., Chen, S., Zhang, Z., Liu, Y., Montardy, Q., Tang, Y., Wei, P., Liu, N., Li, L., Song, R., Lai, J., He, X., Chen, C., Bi, G., Feng, G., Xu, F., & Wang, L. (2019). A VTA GABAergic Neural Circuit Mediates Visually Evoked Innate Defensive Responses. Neuron, 103(3), 473–488.e6. https://doi.org/10.1016/j.neuron.2019.05.027

